# Population genomic evidence of selection on structural variants in a natural hybrid zone

**DOI:** 10.1101/2022.01.14.476419

**Authors:** Linyi Zhang, Samridhi Chaturvedi, Chris C. Nice, Lauren K. Lucas, Zachariah Gompert

## Abstract

Structural variants (SVs) can promote speciation by directly causing reproductive isolation or by suppressing recombination across large genomic regions. Whereas examples of each mechanism have been documented, systematic tests of the role of SVs in speciation are lacking. Here, we take advantage of long-read (Oxford nanopore) whole-genome sequencing and a hybrid zone between two *Lycaeides* butterfly taxa (*L. melissa* and Jackson Hole *Lycaeides*) to comprehensively evaluate genome-wide patterns of introgression for SVs and relate these patterns to hypotheses about speciation. We found >100,000 SVs segregating within or between the two hybridizing species. SVs and SNPs exhibited similar levels of genetic differentiation between species, with the exception of inversions, which were more differentiated. We detected credible variation in patterns of introgression among SV loci in the hybrid zone, with 562 of 1419 ancestry-informative SVs exhibiting genomic clines that deviating from null expectations based on genome-average ancestry. Overall, hybrids exhibited a directional shift towards Jackson Hole *Lycaeides* ancestry at SV loci, consistent with the hypothesis that these loci experienced more selection on average then SNP loci. Surprisingly, we found that deletions, rather than inversions, showed the highest skew towards excess introgression from Jackson Hole *Lycaeides.* Excess Jackson Hole *Lycaeides* ancestry in hybrids was also especially pronounced for Z-linked SVs and inversions containing many genes. In conclusion, our results show that SVs are ubiquitous and suggest that SVs in general, but especially deletions, might contribute disproportionately to hybrid fitness and thus (partial) reproductive isolation.

## Introduction

Understanding the causes and evolution of species boundaries remains a central focus in evolutionary biology (Barton and Bengtsson 1986; Harrison and Larson 2014; Seehausen et al. 2014; Nosil et al. 2017, 2021). In particular, how species boundaries are maintained with ongoing gene flow is debated (Endler 1977; Felsenstein 1981; Servedio et al. 2011; Smadja and Butlin 2011; Nosil 2012). One possibility is that structural DNA changes, that is, genome rearrangements including deletions, duplications, insertions, translocations and inversions, promote speciation and maintain species boundaries despite opportunities for intermittent or ongoing gene flow (reviewed in Wright 1978; Zhang et al. 2021a). Several mechanisms exist whereby structural changes to the genome might drive speciation. For example, structural genome rearrangements can cause meiosis to fail in hybrids leading to hybrid sterility, a common and effective barrier to gene flow (Stebbins 1958; Wright 1978; Coyne and Orr 2004; Zanders et al. 2014). Structural variants (SVs) can also directly cause reproductive isolation through their effects on traits, if, for example, insertions or deletions involve functionally important genes, or if the breakpoints of SVs disrupt a reading frame or alter gene expression (Lynch et al. 2001; Serrato-Capuchina and Matute 2018; Villoutreix et al. 2020; Zhang et al. 2021a). For instance, the insertion of a large transposable element altered the expression of nearby genes in the peppered moth (*Biston betularia*), resulting in the well-known melanic phenotype (Hof et al. 2016).

Beyond these direct effects of SVs, inversions in particular are hypothesized to promote speciation with gene flow by inhibiting recombination within the inverted genomic region (Hoffmann and Rieseberg 2008; Kirkpatrick 2010; Villoutreix et al. 2020; Zhang et al. 2021a). If multiple mutations that contribute directly to reproductive isolation exist, are bought together, or arise within the inverted genomic region, these barrier loci will be inherited as a unit (e.g., as a supergene) and protected from recombination making them more resistant to the homogenizing effects of gene flow (Felsenstein 1981; Noor et al. 2001; Rieseberg 2001; Yeaman 2013). Large inversions could be especially likely to contribute to reproductive isolation via suppressing recombination because their size increases the chance that they will contain multiple, functionally important genes, and thus that they will drive an increase in linkage disequilibrium among a set of physically linked barrier loci. Many such inversion-associated supergenes have been suggested with compelling evidence for a subset, but few have been clearly identified and linked directly to reproductive isolation through their effect on recombination (but see Noor et al. 2001; Kozak et al. 2017).

Indeed, there is increasing evidence of SVs associated with large phenotypic differences in ecologically important traits, including those between ecotypes of sunflowers (Huang et al. 2020), phenological shifts in *Rhagoletis* fruit flies (Feder et al. 2003), the repeated evolution of marine and freshwater ecotypes of three-spined sticklebacks (Jones et al. 2012), and cryptic color pattern variation in *Timema* stick insects (Villoutreix et al. 2020). However, not all of these differences contribute to reproductive isolation (e.g., Comeault et al. 2015), nor is it always clear that the SV mutations directly caused the trait differences. Thus, we lack a general understanding of how often and in what ways SVs contribute to adaptation and speciation. One reason for this lingering uncertainty is that most genomic studies of speciation rely on short-read next-generation sequencing (NGS) (e.g., Illumina sequencing), which has a limited ability to identify and genotype SVs, especially when SV breakpoints are located within repetitive regions in the genome (Mahmoud et al. 2019; Ho et al. 2020). Recent advances in long-read sequencing technology, such as Pacbio SMRT sequencing and Oxford Nanopore sequencing, which routinely produce sequencing fragments of >10 kilobases (kbs), have enhanced our ability to detect SVs (van Dijk et al. 2018; Amarasinghe et al. 2020).

Hybrid zones have long-been recognized as natural laboratories for studying speciation (Hewitt 1988) and thus genomic analyses of hybrid zones with long-read DNA sequence data could be especially informative about the contribution of SVs to speciation. Specifically, gene flow and recombination in hybrid zones create combinations of parental alleles that are then tested by selection under natural conditions (Rieseberg 2001; Gompert et al. 2012a; Schumer et al. 2018). Patterns of ancestry and introgression in hybrid zones capture the outcome of these processes and can be informative about the overall barrier to gene flow and the contribution of individual loci to this barrier (Barton and Hewitt 1985; Barton and Gale 1993; Gompert et al. 2017). Consequently, genomic studies of hybrid zones have contributed substantially to our knowledge of speciation, by for example, providing evidence that sex chromosomes play a major role in speciation (Payseur et al. 2004; Carling and Brumfield 2008; Chaturvedi et al. 2020) and by identifying genetic regions with restricted or aberrant patterns of introgression suggestive of barrier loci (i.e., speciation genes) (e.g., Teeter et al. 2010; Parchman et al. 2013; Knief et al. 2019; Wagner et al. 2020; Semenov et al. 2021). Long-read DNA sequencing should allow more comprehensive and accurate genotyping of SVs across hybrid zones (van Dijk et al. 2018; Amarasinghe et al. 2020), and thus make it possible to study patterns of ancestry and introgression for SVs in hybrids at a genomic scale. Doing so should allow us to better test long-standing theoretical predictions about the mechanisms of SVs in promoting reproductive isolation (Weissensteiner et al. 2020). For example, we can ask whether inversions show reduced or otherwise aberrant introgression compared to other loci (including other types of SVs) as predicted from their role in recombination suppression, or whether SVs in general show evidence of greater selection in hybrids than point mutations, as predicted if SVs often act as major effect mutations by deleting, disrupting or otherwise altering genes and gene expression.

In this study, we used Oxford nanopore long-read DNA sequencing to analyze patterns of introgression for SVs in a hybrid zone between two *Lycaeides* butterfly lineages. *Lycaeides* consists of a complex of multiple species and hybrid lineages of small blue butterflies in North America (Gompert et al. 2014). Hybridization between *Lycaeides idas* and *L. melissa* occurred in the central Rocky Mountains ~14,000 years ago (following the retreat of Pleistocene glaciers) and has resulted in a series of admixed, partially stabilized populations in Jackson Hole valley and the surrounding mountains (hereafter, Jackson Hole *Lycaeides*) (Gompert et al. 2010, 2012a, 2014). In the last two hundred years, these ancient hybrids have secondarily hybridized with *L. melissa*, creating a narrow (1-2 km) contemporary hybrid zone near Dubois, Wyoming, USA. In this study, we focus on this contemporary hybrid zone. Previous work based on SNP data showed that these contemporary hybrids exhibit a wide range of hybrid indexes but that they have, on average, inherited more of their genomes from Jackson Hole *Lycaeides* than *L. melissa*, with a more pronounced excess of Jackson Hole *Lycaeides* ancestry on the Z sex chromosome (Chaturvedi et al. 2020).

In the present study, we first describe the number, size, type, and location of SVs across the genome. We then quantify patterns of population genetic differentiation between *L. melissa* and Jackson Hole *Lycaeides* at these SV loci. Next, we use genomic cline analysis to quantify patterns of introgression within the Dubois hybrid zone for a set of 1419 ancestry informative SVs. We then address the following questions based on analyses of these introgression patterns: 1) Is there evidence that SVs experience more selection on average in hybrids than point mutations (SNPs)? 2) Is the Z sex chromosome enriched for SV loci with patterns of introgression that deviate from null expectations (as was the case for SNPs in Chaturvedi et al. 2020)? 3) Do patterns of introgression differ for different types of SVs, and do inversions specifically show the largest deviations from null expectations? 4) Are patterns of introgression predicted by SV size or by the number of genes within an SV? 5) How well does locus-specific differentiation between parental taxa predict patterns of SV introgression in the hybrid zone?

## Methods

### Study system

*Lycaeides idas* and *L. melissa* diverged from an ancestral species that colonized North America from Asia about 2.4 million years ago (Gompert et al. 2008; Vila et al. 2011). At present, *L. idas* butterflies occur in northwestern North America including much of Canada, the northwestern USA and the northern and central Rocky Mountains, whereas *L. melissa* are found throughout the western USA. The two species’ ranges thus overlap in parts of the Rocky Mountains and northwestern USA (Scott 1992; Gompert et al. 2008). *Lycaeides idas* and *L. melissa* differ in male genitalic morphology, wing pattern, host-plant use, and phenology (Gompert et al. 2010, 2013a,b; Lucas et al. 2018). Microhabitats where two species occur also tend to differ as *L. melissa* more often occupy dry high desert, grassland, or agricultural areas, whereas *L. idas* are often found in more mesic montane environments (Scott 1986). These two species came into secondary contact in the central Rocky Mountains within the past 14,000 years where they hybridized resulting in a broad, patchy hybrid zone comprising a series partially stabilized ancient, admixed populations in the Jackson Hole valley, the Gros Ventre Mountains and the Yellowstone Plateau of northwestern Wyoming (USA) (Gompert et al. 2010, 2012a, 2014). These admixed populations, (i.e., Jackson Hole *Lycaeides*), carry more ancestry from *L. idas* than *L. melissa*, consistent with a tendency for selection to favor *L. idas* alleles in the environment where these hybrids occur, which contains the same host plant used by nearby *L. idas* populations and is generally more *L. idas-like* (Gompert et al. 2012a).

Within the past two hundred years, these ancient hybrid populations have come into contact with and hybridized with *L. melissa* found on feral, roadside alfalfa *(Medicago sativa)* near the town of Dubois, Wyoming (USA), creating a narrow (1-2 km) contemporary hybrid zone (Chaturvedi et al. 2020). The hybrid zone lacks a clear spatial gradient instead comprising a single admixed population with parents and many hybrids (Chaturvedi et al. 2020). In this study, we analyze patterns of hybridization and introgression (i.e., the movement of alleles from one genomic background to another) in this contemporary hybrid zone between Jackson Hole *Lycaeides* and *L. melissa.* The high elevation (2115 meters), montane environment here is more similar to locations where *L. idas* and Jackson Hole *Lycaeides* are found, whereas the immediate local habitat borders agricultural fields similar to many *L. melissa* locations and the the main host plant here, alfalfa, is used by *L. melissa*, but not *L. idas* or Jackson Hole *Lycaeides* (Gompert et al. 2013b; Chaturvedi et al. 2020).

### DNA extraction, library preparation and MinION nanopore sequencing

Genomic DNA from a total of 39 male butterflies (males are the homogametic sex), including seven Jackson Hole *Lycaeides*, six *L. melissa*, and 26 *Lycaeides* from the Dubois hybrid zone (Table S1) was isolated with Qiagen’s MagAttract High-Molecular-Weight (HMW) DNA extraction kit (cat. No. 67563; Qiagen Inc., Valencia, CA, USA). The concentration of DNA was assessed using the dsDNA HS assay on a Qubit 4 fluorometer (Thermo Fisher).

We then created a DNA library for each butterfly for whole-genome long-read sequencing on the Oxford MinION platform. Specifically, we first repaired DNA molecules with the NEBNext FFPE DNA Repair Mix (NEB M6630) and NEBNext Ultra II end repair/dA-tailing Module (NEB E7546). This was done using 0.5-1.0 ug genomic DNA per sample and in accordance with the manufacturer’s protocol. Individuals with similar DNA concentrations were barcoded using the Native Barcoding Expansion Pack 1-12 (EXP-NBD 104) and subsequently pooled to reach a total amount of 1 μg of genomic DNA or a total number of 12 individuals (whichever came first); the number of individuals per pooled library was between 4 and 12 (Table S2). We then ligated adaptor oligos for sequencing onto the barcoded fragments in each pooled library using the Ligation Sequencing kit (SQK-LSK109). The ligation product from each library was then loaded onto R9.4 flow cell (FLO-Min106, ONT) for a sequencing run of a full 72 hours on a MinION. This generated an average of 7 GB of sequence data per library, with a N50 read length of 1114 to 20,944 bps (mean read length = 9073 bp; maximum read length of 342,840 bp).

### Base calling, structural variant calling, and variant filtering

Base calling was performed using the ont-guppy algorithm from guppy_basecaller (version 4.2.2). We then used guppy_barcoder (version 4.2.2) to demultiplex the nanopore sequence reads and subsequently remove the barcode sequences. Next, SVs, including inversions, tandem duplications, insertions and deletions, relative to the newly updated *L. melissa* genome (see supplemental material) were identified following Oxford Nanopore’s suggested best practices (see https://github.com/nanoporetech/pipeline-structural-variation) (Jain et al. 2018; Jiang et al. 2020; Ren and Chaisson 2021). Here, a duplication is a specific type of insertion where the inserted segment is a tandem repeat of the existing DNA region, and in general insertions versus deletions are defined as added or missing DNA bases relative to the reference genome. Specifically, we used Minimap2 (version 2.17) to align the nanopore reads to the reference genome with default parameters including a mismatch penalty set at 4 and gap open penalty at 24 (Li 2018). We then used Sniffles (version 1.0.12) with default settings to identify SVs (Sedlazeck et al. 2018). This was done separately for each individual. Sniffles detects SVs from long-read alignments by using both within-alignment and split-read information, as small indels can be spanned within a single alignment, whereas large SVs lead to split-read alignment (Sedlazeck et al., 2018). Importantly, Sniffles filters false SV signals by considering both minimum read support as well as consistency of the breakpoint position and size. Because loci with excessive htererozygosity are likely caused by erroneous mapping of transposable elements to the reference genome (Jaegle et al. 2021), we excluded SVs with excessive htererozygosity, defined for this purpose as all individuals in either species initially being called heterozygotes because they had reads supporting the reference and SV allele. Additionally, we excluded SVs with sequence data for less than 20% of the individuals, which left a total of 290,276 SVs for downstream analysis. Next, we merged SVs across samples using SURVIVOR (https://github.com/fritzsedlazeck/SURVIVOR; version 1.0.3) (Jeffares et al. 2017). When merging, we only considered SVs of at least 30 bps and merged those that overlapped with a maximum allowed distance of 1 kb as measured pairwise between start and stop coordinates.

SURVIVOR does not compute genotype likelihoods, but rather only reports the number of reads supporting the reference and alternative alleles and a called genotype based on a hard-threshold number of reads (Jeffares et al. 2017). This ignores uncertainty in SV genotype. Thus, rather than using the genotype calls, for each individual and locus we computed genotype likelihoods for being homozygous for the reference allele, homozygous for the structural variant allele or heterozygous. This allowed us to propagate uncertainty in structural variant genotypes to downstream analyses (as is more routinely done with SNP data sets, e.g., Gompert et al. 2014). Specifically, the genotype likelihoods were calculated based on the number of reads supporting the reference and SV alleles and an assumed per-read SV error rate:

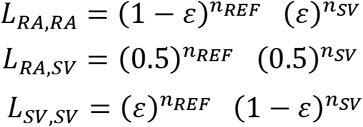

Here, *L_RA,RA_, L_RA,SV_* and *L_SV,SV_* refer to the relative likelihoods of being homozygous for the reference allele, heterozygous and homozygous for the SV allele, respectively. The number of reads supporting the reference and structural variant allele for an individual are denoted *nREF* and *nSV* and *ε* refers to the error rate. This is not equivalent to the probability of a single base being erroneous, but rather the probability of a sequence read erroneously supporting the reference or structural variant allele. Because this error rate is not precisely known, we focus our main results on the case of *ε* = 0.01, but show results based on *ε* = 0.05 in the online supplement (both gave similar results; see the Supplemental Material for details).

With the computed genotype likelihood, allele frequencies for each SVs in *L. melissa*, Jackson Hole *Lycaeides* and the Dubois hybrid zone were estimated using estpEM (version 0.1) (Sorria-Carrasco et al. 2014; DRYAD https://doi.org/10.5061/dryad.nq67q) with a convergence tolerance of 0.001 and 20 EM iterations. The estpEM program infers allele frequencies via an expectation-maximization (EM) algorithm while accounting for the uncertainty in genotypes (Li 2011). Only SVs with a minimum (global) minor allele frequency of 0.05 computed from all individuals were retained. This gave us a total 127,574 SVs. For downstream population introgression analyses (see below), we analyzed only SVs > 1000 bps in length, which left us with 15,319 SVs.

### Validating the structural variants with additional whole-genome sequencing

We generated additional whole-genome sequence data to validate the SVs from the nanopore data. Specifically, we wanted to know whether a non-trivial subset of the SVs identified by nanopore sequencing overlapped with those found by traditional, mate-pair Illumina sequencing. For this, we used genomic data that were generated as part of a larger effort to produce reference genomes for multiple *Lycaeides* species, but that have not been previously published. Thus, for this data set we focused on a smaller number of individuals, one from each of five nominal lineages of *Lycaeides* in North America: *L. idas* (from Trout Lake, WY), *L. melissa* (from Bonneville Shoreline Trail, UT), *L. anna* (from Yuba Gap, CA) and admixed lineages in the Sierra Nevada (from Carson Pass, CA) and Warner mountains (from Buck Mountain, CA) in the western USA. Genomic DNA was isolated from each butterfly using Qiagen’s MatAttract HMW DNA extraction kit (Qiagen, Inc.). We then outsourced library construction and sequencing to Macrogen Inc. (Seoul, South Korea). A 3-kb mate-pair library was created for each butterfly species using the Nextera Mate-pair Library Gel-plus kit (Illumina, Inc.). The libraries were then sequenced on a HiSeq 2000 (2×100 bp reads) generating 669,157,572 Q30 reads (67,584,914,723 bps total or ~30X coverage per genome). SVs were identified by first using the SV callers DELLY and LUMPY and then combining the output from these callers with SURVIVOR (version 1.0.7) (Jeffares et al. 2017) (see the Supplemental Materials for details regarding DNA sequence alignment, SV calling and filtering with the mate-pair data). After filtering, this left us with 12,265 SVs from the mate-pair data DNA sequence data set, including 3058 SVs that were > 1 kbp in length.

### Describing population genetic variation for structural variants in the nanopore data set

With allele frequency estimates using estpEM program, we computed pairwise FST among *L. melissa*, Jackson Hole *Lycaeides* and the Dubois hybrid zone, with each of these treated as a population. F_ST_ was calculated as 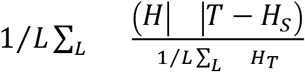, where *H_S_* and *H_T_* are the expected subpopulation and total heterozygosities for each locus. We estimated FST separately for each pair of entities and SV type. Next, we designated a set of ancestry informative SVs as those having an allele frequency difference ≥0.3 between *L. melissa* and Jackson Hole *Lycaeides*, which we used for all subsequent analyses. A total of 1419 SVs met this criterion. We used an admixture model, as implemented in Entropy (version 1.2), to estimate admixture proportions and genotypes for these ancestry informative SVs (Gompert et al. 2014; Shastry et al. 2021). This model is similar to the admixture model in STRUCTURE (Pritchard et al. 2000), except that the uncertainty in genotypes due to limited sequencing coverage and errors are incorporated into the estimation of genotypes and admixture proportions. We fit this model based on our computed genotype likelihoods and assumed two source populations (i.e., *K* = 2 as we had two hybridizing taxa). We ran five Markov chain Monte Carlo (MCMC) chains with k=2, each with 10,000 MCMC iterations, a burn-in of 5000 iterations and a thinning interval of 5. We obtained Bayesian genotype estimates by taking the posterior mean of the number of SV alleles (0, 1, or 2) for each locus (this mean is not constrained to be an integer value). Lastly, we visualized patterns of genetic variation at the ancestry informative SVs using principal component analysis (PCA); this was done in R version 4.0.2 on the centered but not standardized genotype matrix.

### Estimating genomic cline parameters in the Dubois hybrid zone

We quantified patterns of introgression for the SVs in the Dubois hybrid zone to test for differential introgression of SVs relative to each other and relative to average introgression as measured by previously published ancestry informative SNPs. We did this by fitting Bayesian genomic clines with bgc (version 1.05a; https://github.com/zgompert/BGC-Bayesian-genomic-clines) (Gompert and Buerkle 2011, 2012). Using genomic clines, it is possible to quantify introgression into alternative genomic backgrounds even when hybrid zones do not display a strong spatial axis (Gompert and Buerkle 2009, 2011), as is the case for the Dubois hybrid zone. More specifically, deviations between genome-average introgression and the degree of introgression for each SV locus were measured with two cline parameters, *α* and *β*. Cline parameter *α* describes an overall deviation in the probability of ancestry relative to null expectations from an individual’s hybrid index, where a positive (negative) value of *α* indicates an increase (decrease) in the probability of reference species 1 ancestry (*L. melissa* ancestry in our analysis). Cline parameter *β* describes a deviation in the average rate of transition from reference species 0 ancestry to reference species 1 ancestry along the genome-wide admixture gradient. Positive values for *β* indicate a decrease in the rate of transition (i.e., alleles confined mostly to one genetic background), whereas negative values for *β* indicate a decrease in the rate of transition (i.e., alleles from both species flowing more into alternative genomic backgrounds). Selection in hybrids can lead to departures of cline parameters from genome-wide average admixture (Gompert et al. 2012b). For example, selection generated by negative epistatic interactions (DMIs) can cause reduced introgression (increased *β*) or a shift in ancestry towards one of the two species (high negative or positive *α*) (Gompert et al. 2012b). Drift can also cause deviations from genome-wide admixture, but loci with credible deviations for *α* or *β* should be enriched for loci affected (directly or indirectly via LD) by selection (Gompert et al. 2012b). As this concerns selection in hybrids, this includes possible barrier loci contributing to reproductive isolation.

In this study, we were interested in whether patterns of introgression for SVs differed overall from genome-average admixture as measured by non-SV loci (i.e., SNPs). Therefore, we used the SNP data to estimate hybrid index (i.e., admixture proportions) and thus the average extent of introgression, and then fit genomic clines for the SV loci conditional on the SNP estimates of hybrid index. The SNP data were taken from a previous study that included many of the individuals used in this study (Chaturvedi et al. 2020) (see Table S2), with a total of 1164 ancestry-informative markers (allele frequency differences between two parental species ≥0.3). Clines were inferred using a modified version of the bgc software that allows separate sets of loci to be used for fitting clines and estimating hybrid indexes, and that does not require the cline parameters to sum to zero across loci (bgc version 1.05a). Thus, positive *α* values indicate an increase in the probability of *L. melissa* ancestry relative to genome-wide average introgression based on SNPs (and vice a versa for negative α); positive *β* for a structural variant locus would likewise indicate reduced introgression compared to the average for the SNP loci. Genomic clines were fit using MCMC, with five chains each with 200,000 iterations, a 20000-iteration burn-in, and a thinning interval of 5. We inspected the MCMC output to assess convergence of chains to the stationary distribution and combined the output of the five chains. We designated SV loci as credibly deviating from genome-average introgression when the 95% equal-tail probability intervals for α or β excluded 0. Five hundred sixty-two SVs exhibited credible deviation based on α, but none did based on β; we thus focus on α in the tests below.

We next conducted a series of statistical tests to analyze the estimated genomic clines with respect to expectations from the hypothesis that SVs are a key driver of speciation. First, we tested for an overall shift in the direction of introgression for SVs relative to SNPs by estimating the mean value of the cline parameter α across SVs and by determining whether the observed asymmetry in credible positive versus negative α values was consistent with binomial expectations (with probability parameter *p* = 0.5). Then, to determine whether patterns of introgression differed between the autosomes and the Z-chromosome, we compared the mean signed *α* cline parameter for Z-linked SVs to null expectations generated by randomly sampling the same number of SV loci from the autosomes; this was done 1000 times. Next, to test whether patterns of introgression differed for different types of SVs, and specifically whether inversions showed the largest deviations from null expectations, we compared the mean value of signed *α for* inversions to null expectations generated by a randomly sampling a subset of the same number of loci from all other SV types (this was also done 1000 times).

We then tested the hypothesis that SV length or the number of genes in a SV affects patterns of genetic differentiation between species and patterns of introgression in the hybrid zone (gene numbers were taken from the genome annotation described in Chaturvedi et al. 2020 and then transferred to our updated *L. melissa* genome; see the Supplemental Materials for details). As expected, SV length and number of genes within a SV were positively correlated (Pearson r = 0.863, 95% CI: 0.862-0.865, P <0.001), and thus we fit multiple regression models with both covariates simultaneously to better assess their marginal effects. Specifically, we fit linear models for *α* (one of the cline parameters) or the absolute value of *α* as a function of SV length (in bps), number of annotated genes in each SV, and their interaction for each class of SVs (covariates were centered and standardized prior to analysis). This was done with the lm function in R version 4.0.2. Here, models for signed values of *α* detect factors associated with an overall directional shift (for or against Jackson Hole or *L. melissa* ancestry), whereas models for the absolute value of *α* detect factors associated with the magnitude of introgression independent of the direction. We fit similar multiple regression models for the 1419 ancestry informative SVs for genetic differentiation (F_ST_, logit transformed) between the parental lineages as well.

Finally, we tested for an association between genetic differentiation at individual SV loci between the allopatric Jackson Hole *Lycaeides* and *L. melissa* populations and patterns of introgression for these loci in the hybrid zone. A positive association between F_ST_ and the absolute value of α would be expected if differentiation in allopatry was driven by divergent selection and the same SV loci under divergent selection in allopatry also affected hybrid fitness in the Dubois hybrid zone (e.g., Gompert et al. 2012b). Moreover, a correlation between F_ST_ and the signed (directional) value of *α* might be expected if alleles from one parental species are disproportionately favored in the hybrid zone, either because of environment-dependent fitness or because of DMIs causing selection against minor parent ancestry (e.g., Schummer et al. 2018; Chaturvedi et al. 2020). Thus, Pearson correlation tests were conducted for FST versus both absolute *α* and signed *α* (this was done in R version 4.0.2). Importantly, whereas ancestry informativeness can create spurious correlations between FST and cline parameters, past work suggests this is unlikely when considering only ancestry informative markers as was done here (Gompert et al. 2012b).

## Results

### Number, length, and distribution of SVs across genome

Using long-read nanopore sequencing, we identified a total of 127,574 SVs that covered 30.4% of the genome, including 15,319 SVs >1000 bp in length. Among all SVs identified, there were 68,927 deletions covering 18.4% of the genome, 56,527 insertions covering 4.0% of the genome, 1503 inversions covering 7.2% of the genome and 617 duplications covering 0.7% of the genome (Figure 1A,B; see Table 1 and Figure 2A for range of SV sizes). Considering only SVs > 1kbp in length, we identified 13,359 deletions covering 15.8 % of the genome, 1073 inversions covering 7.1% of the genome, 771 insertions covering 1.3% of the genome and 116 duplications covering 0.1% of the genome (Figure 1C, 1D; Table 1).

**Figure 1.**
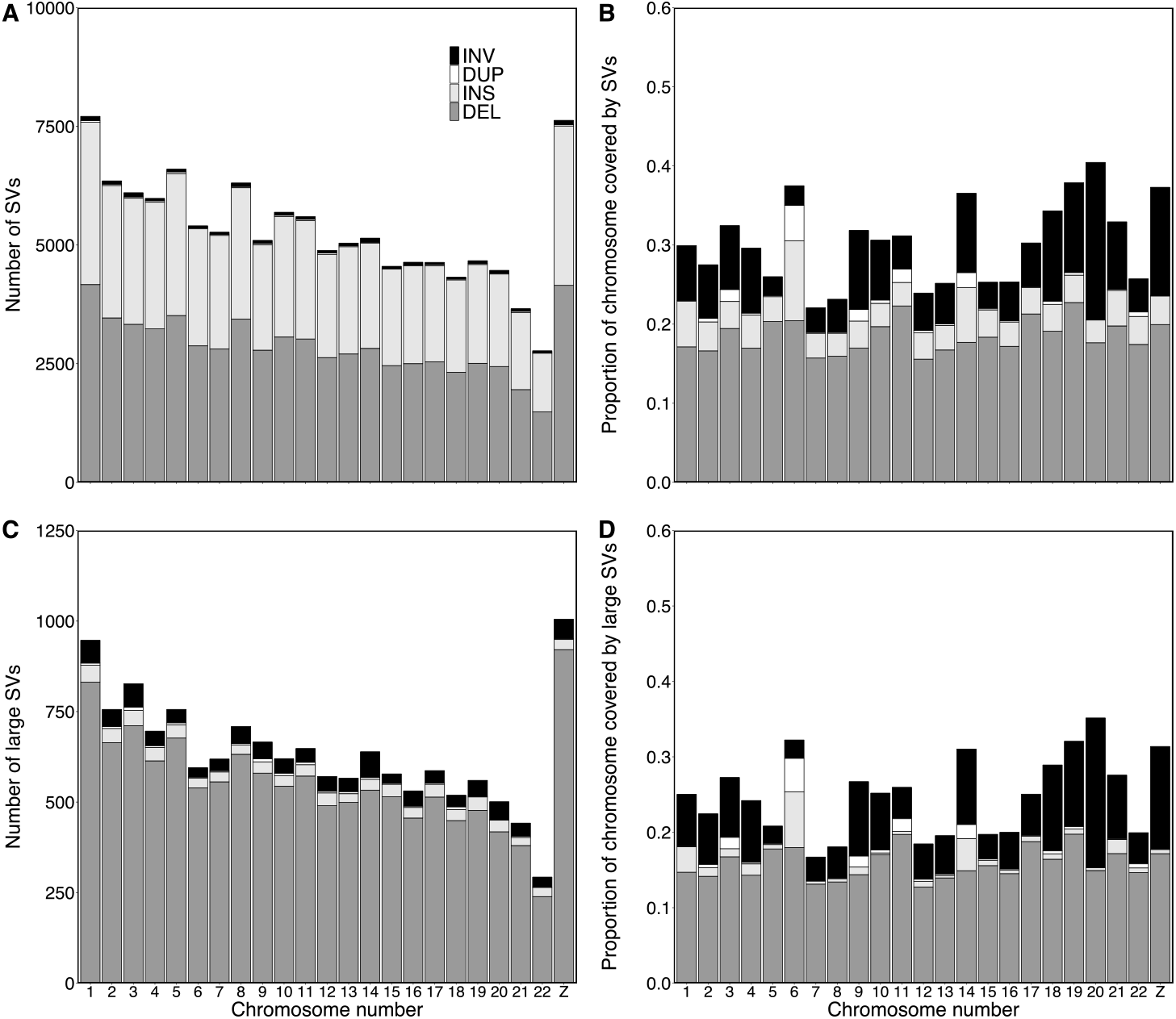
(A) Number of structural variants (SVs) across the genome. (B) Proportion of each chromosome covered by SVs. (C) Number of large SVs (>1000 bp) across the genome. (D) Proportion of each chromosome covered by large SVs (>1000 bp). Bars with different colors denote different SV types, as shown in the legend: INV = inversion, DUP = duplication, INS = insertion, and DEL = deletion.

**Table 1.** Number and length (mean, 2.5th and 97.5th percentiles) of SVs detected by nanopore sequencing; SV types: INV = inversion, DUP = duplication, INS = insertion, and DEL = deletion.

|  | SV type | No. SVs | Length of SVs (bp) |  |  |
| --- | --- | --- | --- | --- | --- |
|  |  |  | Mean | 2.50% | 97.50% |
| **All SVs** | DEL | 68927 | 1571 | 42 | 11554 |
|  | DUP | 617 | 5472 | 51 | 20393 |
|  | INS | 56527 | 359 | 43 | 803 |
|  | INV | 1503 | 41306 | 175 | 422305 |
| **SVs with length>1000 bp** | DEL | 13359 | 7171 | 1035 | 28073 |
|  | DUP | 116 | 27697 | 1074 | 356753 |
|  | INS | 771 | 8751 | 1000 | 42286 |
|  | INV | 1073 | 57670 | 1104 | 542297 |

**Figure 2.**
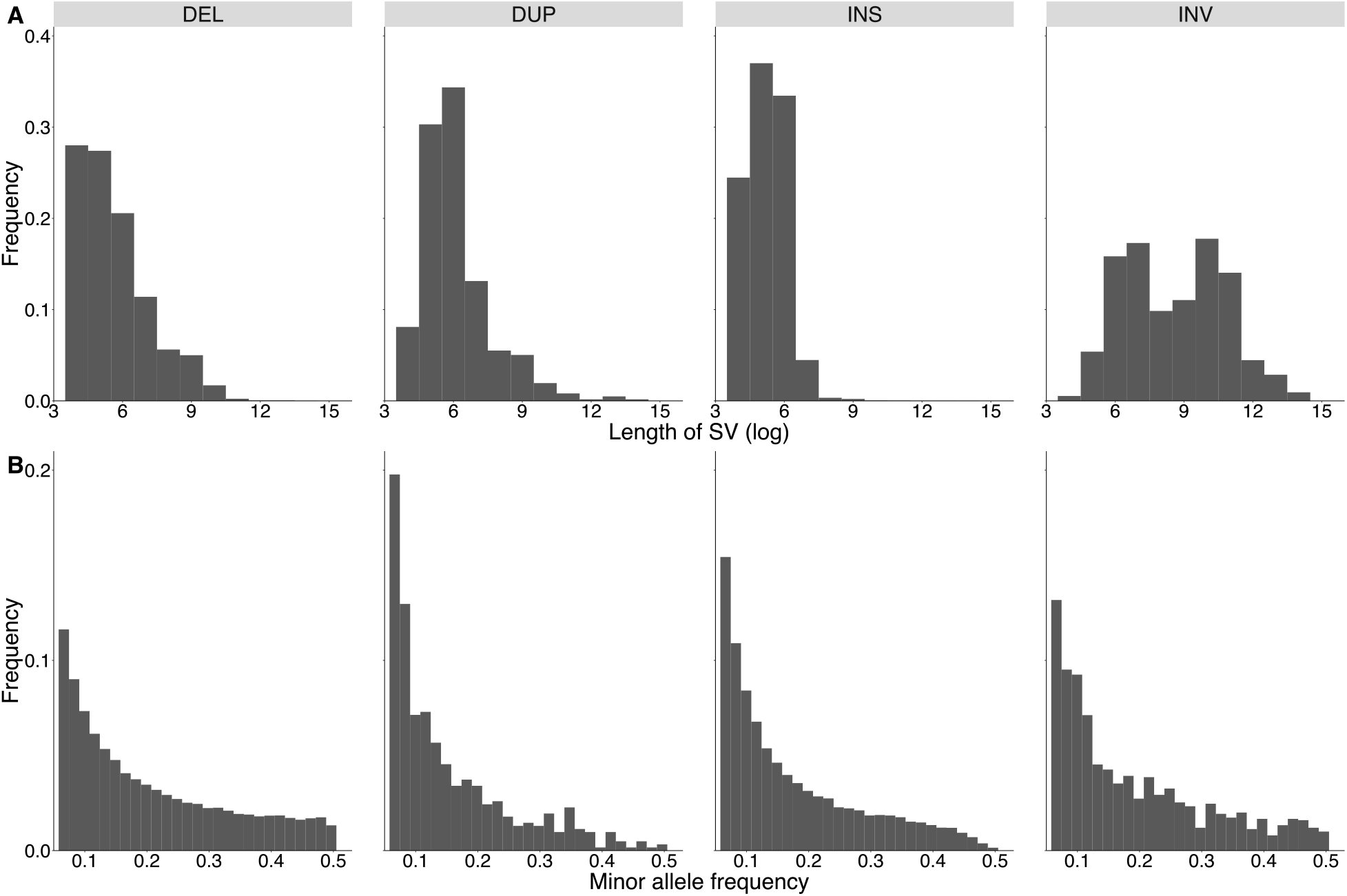
Histograms of distribution of (A) length of SV (log) across different types of structural variants, (B) minor allele frequency of all 39 individuals of *Lycaeides.*

We detected 12,265 SVs using mate-pair sequencing, 32.5% of which corresponded with SVs identified from nanopore sequencing (Table 2). Similar to the nanopore data, most of the SVs detected from the mate-pair data were deletions (11,594 out of 12,265), followed by 224 duplications, and 156 inversions (Figure S2, S3).

**Table 2.**
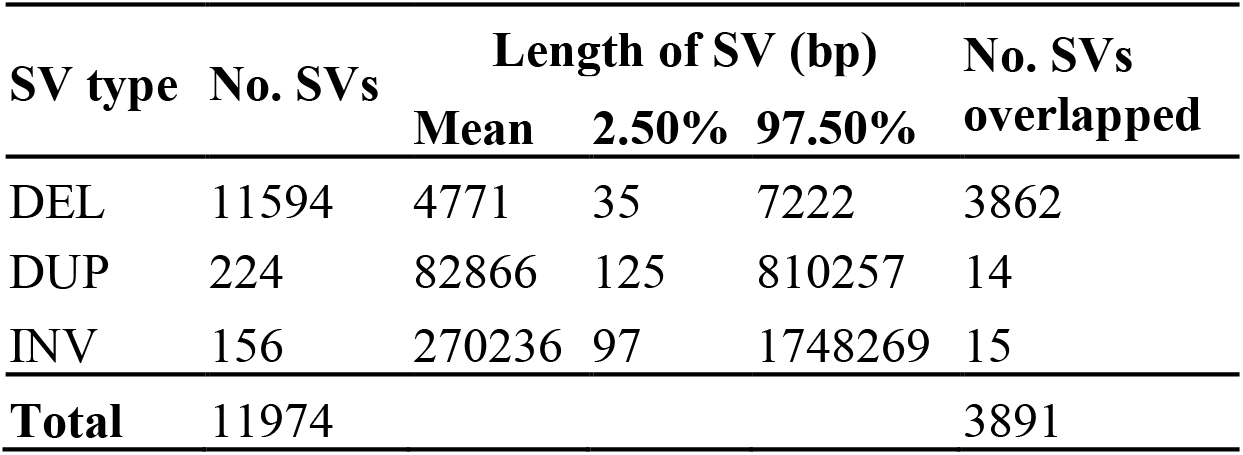
SVs detected by mate-pair sequencing and number of mate-pair SVs that overlapped with those found by nanopore sequencing; SV types: INV = inversion, DUP = duplication, INS = insertion, and DEL = deletion.

### Population genomic patterns of structural variation

The distribution of allele frequencies was skewed towards rare alleles among *Lycaeides* individuals (Figure 2B), which is consistent with recent studies in grapevine, Drosophila, and songbirds SV (Chakraborty et al. 2019; Zhou et al. 2019; Weissensteiner et al. 2020). Most SVs exhibited only subtle allele frequency differences between *L. melissa* and Jackson Hole *Lycaeides* (mean difference = 0.118), but 1419 SVs exhibited allele frequency differences exceeding 0.3, and four SVs (all deletions) were fixed or nearly fixed between these taxa (i.e., allele frequency difference ~1.0) (Figure S1). We detected modest differentiation between Jackson Hole *Lycaeides* and *L. melissa* based on the full set of 127,574 SVs (F_ST_ = 0.048) (Table 3). This is similar to an earlier estimate of genetic differentiation between these lineages based on SNP data (F_ST_ = 0.050) (Chaturvedi et al. 2020). However, genetic differentiation (F_ST_) for inversions was higher than for other SV types or SNPs (F_ST_ = 0.070) (Table 3). Lastly, genetic differentiation was positively associated with deletion length and the number of genes in a deletion (with a positive interaction between these two covariates) (Table 4), but no such associations were detected for the other SV types.

**Table 3.**
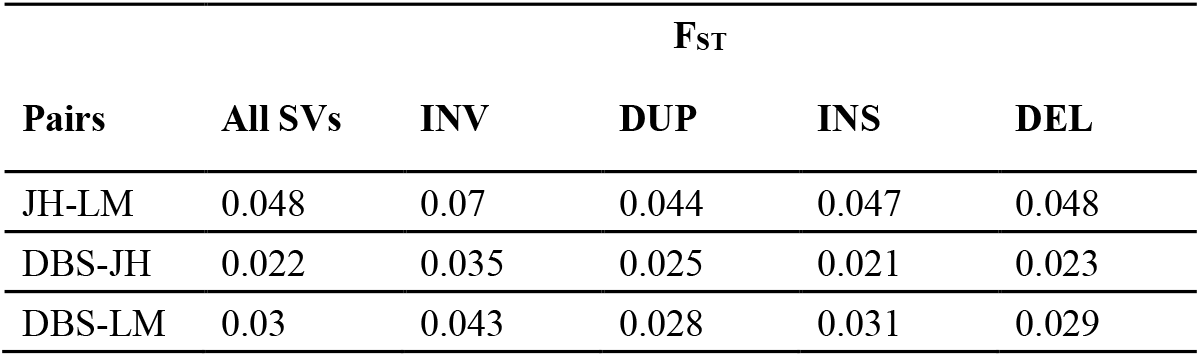
Comparison of pairwise F_ST_ across different structural variant types. Lineage codes: JH = Jackson Hole *Lycaeides*, LM = *L. melissa*, DBS = Dubois hybrid zone. SV types: INV = inversion, DUP = duplication, INS = insertion, and DEL = deletion.

| Pairs | $F_{ST}$ | | | | |
| --- | --- | --- | --- | --- | --- |
|  | All SVs | INV | DUP | INS | DEL |
| JH-LM | 0.048 | 0.07 | 0.044 | 0.047 | 0.048 |
| DBS-JH | 0.022 | 0.035 | 0.025 | 0.021 | 0.023 |
| DBS-LM | 0.03 | 0.043 | 0.028 | 0.031 | 0.029 |

**Table 4.** Results of multiple linear regression models of F_ST_ against length and number of genes within each SV. SV types: INV = inversion, DUP = duplication, INS = insertion, and DEL = deletion.

| SV type | Factor | Estimate | SE of estimate | *t*-value | *P*-value |
| --- | --- | --- | --- | --- | --- |
| DEL | length | 0.094 | 0.023 | 4.103 | < **0.001** |
|  | number of genes | 0.055 | 0.024 | 2.242 | **0.025** |
|  | length: no. genes | -0.004 | 0.001 | -5.345 | < **0.001** |
| DUP | length | -0.009 | 0.103 | -0.090 | 0.928 |
|  | number of genes | -0.007 | 0.100 | -0.068 | 0.946 |
|  | length: no. genes | 0.001 | 0.003 | 0.425 | 0.671 |
| INS | length | -0.066 | 0.066 | -0.994 | 0.320 |
|  | number of genes | -0.030 | 0.044 | -0.682 | 0.495 |
|  | length: no. genes | 0.004 | 0.002 | 1.502 | 0.133 |
| INV | length | 0.019 | 0.017 | 1.111 | 0.267 |
|  | number of genes | 0.009 | 0.025 | 0.378 | 0.705 |
|  | length: no. genes | -0.001 | 0.001 | -1.142 | 0.254 |

PCA and admixture proportion estimates from ancestry informative SVs suggested generally similar patterns of genetic variation in the parental lineages and hybrids as past work using SNPs (e.g., Chaturvedi et al. 2020) (Figure 3). For example, genetic PC scores from SNPs and SVs were highly positively correlated (Pearson *r* = 0.772, *P* < 0.0001) and *Lycaeides* from the Dubois hybrid zone displayed a range of admixture proportions (i.e., hybrid indexes) from nearly pure Jackson Hole *Lycaeides* to *L. melissa* with many intermediates (Figure 3B, 3C).

**Figure 3.**
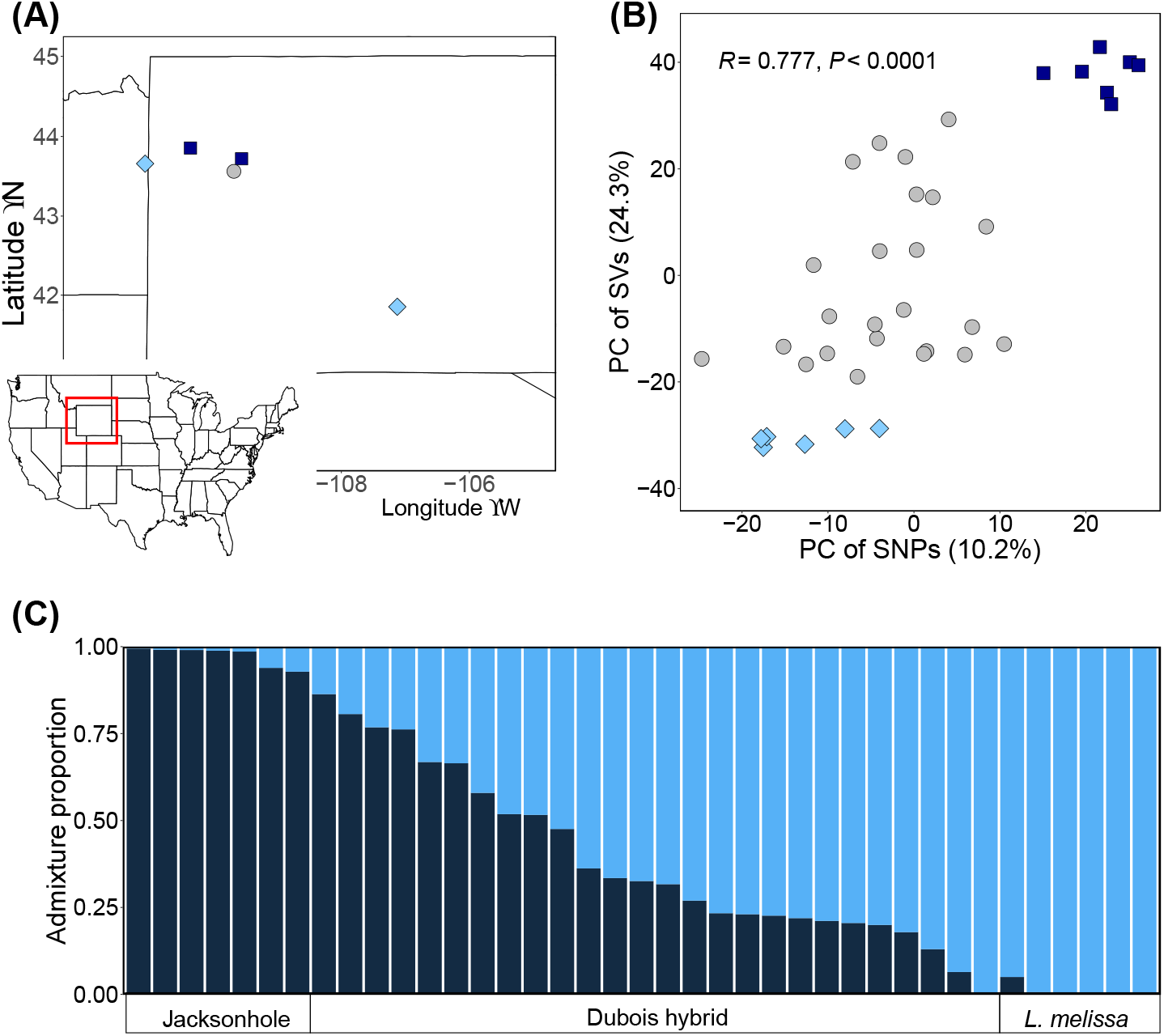
(A) Collection sites, (B) Plot of PC score 1 of large, ancestry informative SVs (>1000bp, allele frequency differences>0.3) against PC score 2 of SNPs, (C) Entropy plot from large, ancestry informative SVs (>1000bp, allele frequency differences>0.3), y axis is the Bayesian estimates of admixture proportion.

### Pattern of introgression of SVs across Dubois hybrid zone

We observed qualitative variation in patterns of introgression in the hybrid zone for the 1419 SVs (Figure 4), which was confirmed quantitatively using genomic cline analysis (Figure 5). Specifically, credible variation in patterns of introgression across the genome were detected for 562 SVs (cases where 95% equal-tail probability intervals [ETPIs] for genomic cline parameters did not overlap with zero) (Figure 5B, C). Relative to genome-average introgression computed from ancestry-informative SNPs, a total of 152 SVs showed credible excess introgression from *L. melissa* (95% ETPIs for cline parameter *α* > 0) and 410 SVs exhibited credible excess introgression from Jackson Hole *Lycaeides* (95% ETPIs for cline parameter *α* < 0). Consequently, relative to average introgression for SNPs, SVs showed an excess of Jackson Hole *Lycaeides* ancestry in the hybrid zone (mean *α* ± SE = −0.295 ± 0.018; probability of getting 410 credible excessive Jackson Hole ancestry loci out of 562 credible excess introgress loci: *P* < 0.0001, Figure 5C). No SVs displayed credible shifts in the rate or extent of introgression (95% ETPIs for cline parameter *β* > 0 or *β* < 0) (detecting credible deviations based on beta often requires larger sample sizes across the range of hybrid indexes).

**Figure 4.**
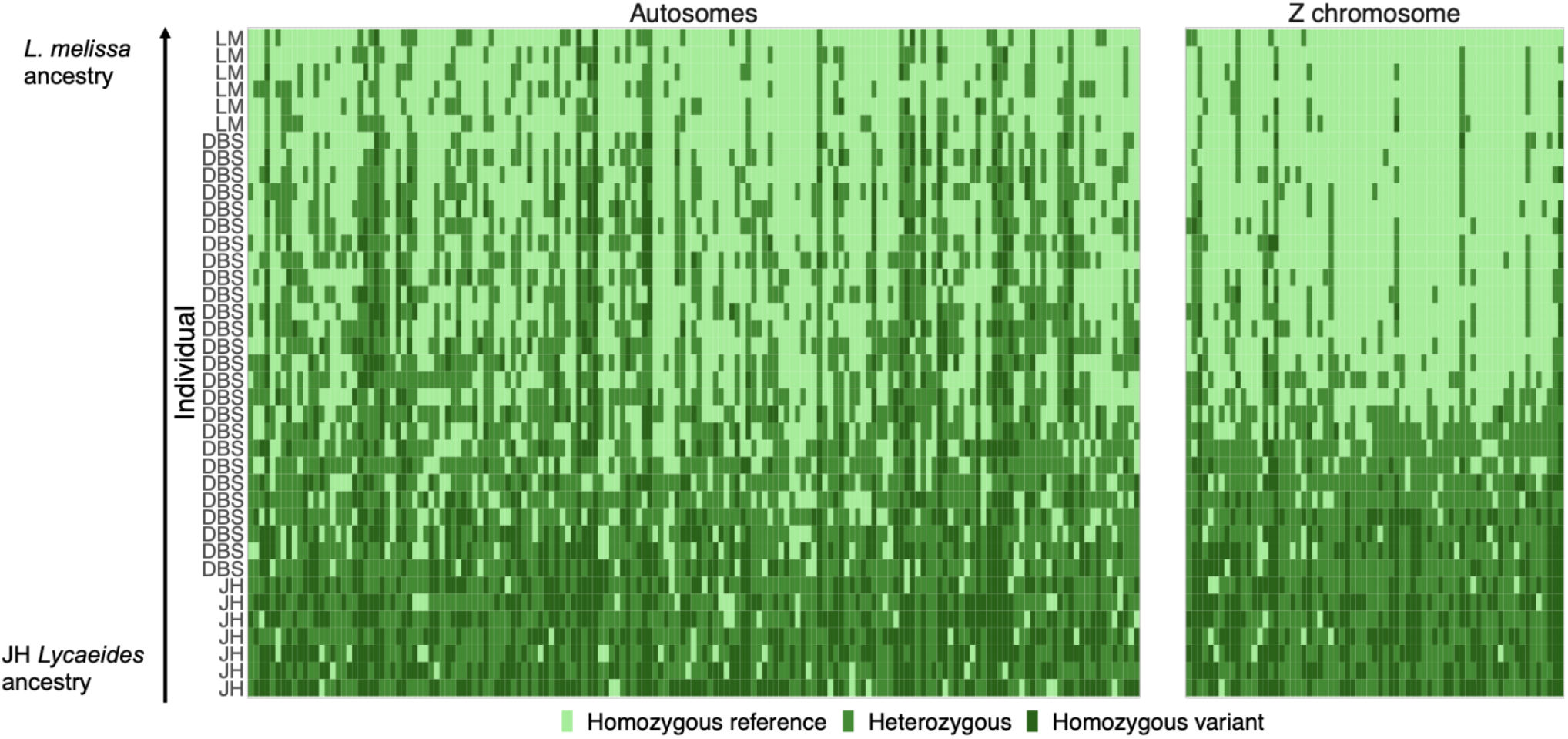
Genotypes of ancestry informative loci (allele frequency differences >0.5) on autosomes vs. the Z chromosome across individual butterflies. Each colored rectangle denotes the genotype estimate for a SV locus (x-axis) and individual (y-axis). Genotype estimates were rounded to the nearest whole number for visualization. Loci on the x-axis are ordered by first chromosome number, then by position of the locus, and lastly by types of SVs and length. Individuals are sorted by the order of the hybrid index (proportion of ancestry from *L. melissa*). JH stands for parental species Jackson Hole *Lycaeides*, DBS stands for Dubois hybrid individuals, LM stands for parental species *L. melissa.*

**Figure 5.**
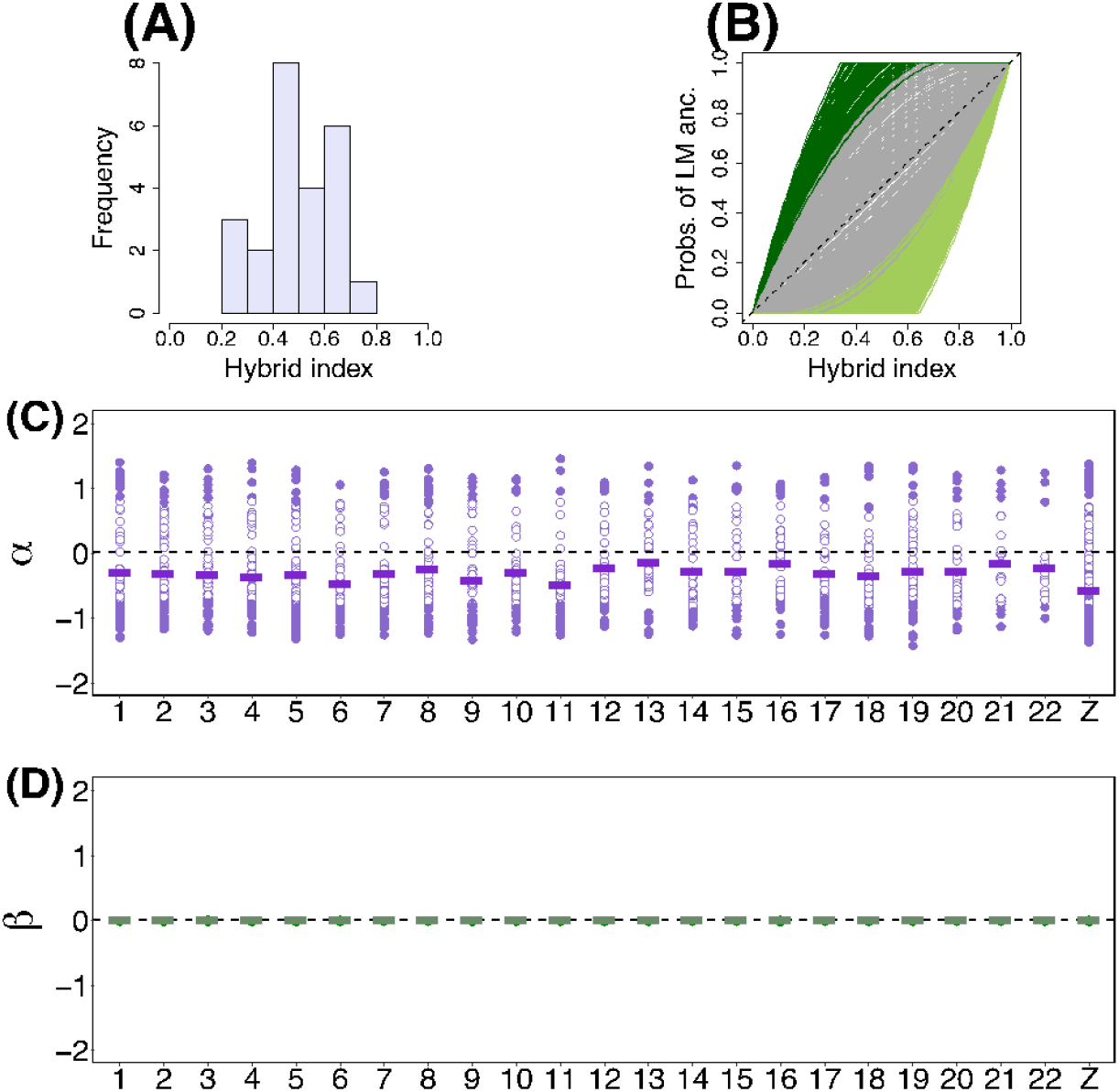
Summary of the genomic cline analysis. (A) Distribution of hybrid index using structural variant data. (B) Estimated genomic cline relative to SNPs using ancestry-informative SVs. Each solid line gives the estimated probability of Jackson Hole (JH) ancestry for a structural variant locus. Green lines denote cases of credible directional introgression (95% ETPIs for α that do not overlap with zero), gray lines denote clines not significantly different from the genome average. The dashed line gives the null expectation based on genome-wide admixture. Distribution of cline parameters *α* (**C**) and *β* (**D**) across different chromosomes. Solid circles indicate loci with cline parameters indicating credible deviations from genome-average introgression (95% ETPIs that do not overlap with zero), whereas unfilled circles indicate loci not significantly different from the genome average. Solid horizontal lines are the mean value of cline parameters across all loci at each chromosome. The dashed horizontal line gives the null expectation based on genome-wide admixture using SNP loci.

Structural variants exhibiting excess directional introgression were distributed across the 23 *Lycaeides* chromosomes, but SVs on the Z-chromosome showed significantly more excess Jackson Hole *Lycaeides* introgression compared to SVs on the autosomes (permutation test with 1000 permutations: mean difference in signed *α* [Z - autosomes] = −0.268, *P* < 0.001). The degree of directional introgression also differed among different types of SVs, with deletions showing significantly more excess Jackson Hole *Lycaeides* introgression than inversions (permutation test with 1000 permutations: mean differences in signed *α* [deletion – inversion] = −0.232, *P* < 0.001, Figure 6A).

**Figure 6.**
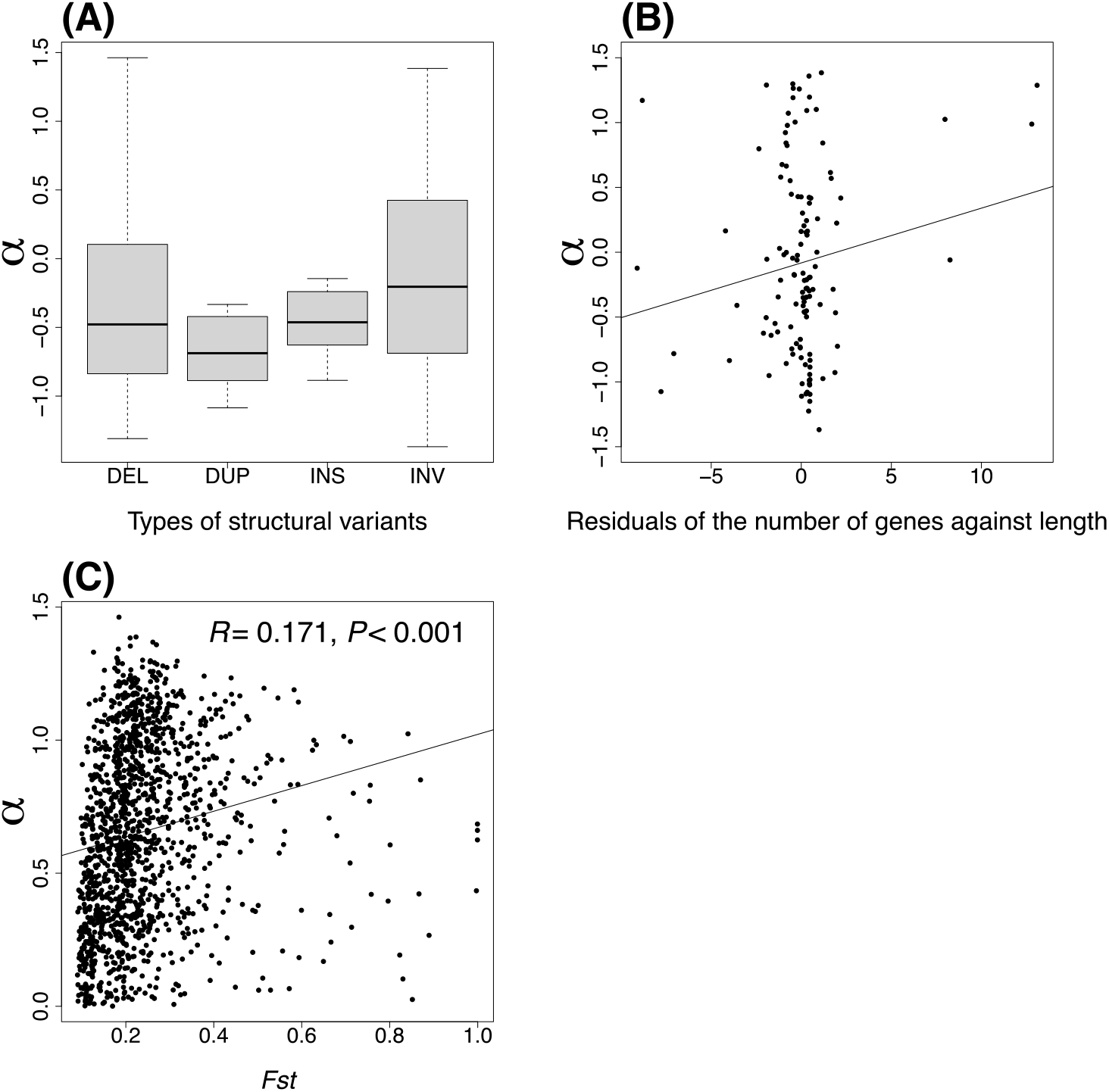
(A) Comparison of cline parameter *α* across different structural variant types. (B) Correlation between *α* and the residuals of the number of genes against the length across all inversions. (C) Correlation between *α* and F_ST_ across loci.

We did not detect significant associations of SV length or the number of genes in SVs with the absolute value of α (i.e., the extent of excess introgression regardless of direction) for any of the SV types based on the multiple regression analyses (Table S3). However, the degree of directional introgression (signed *α*) was positively associated with the number of genes contained (regression coefficient = 0.19, *t* = 2.35, *P* = 0.021) but not SV length for inversions (regression coefficient ≈ 0, *t* = −0.79, *P* = 0.431; Table 5 and Figure 6B). In contrast, the degree of directional introgression (signed *α*) was positively associated with the length (regression coefficient =0.248, *t* = 3.02, *P* = 0.003) but not the number of genes contained within deletions (regression coefficient =0.02, *t* = 0.15, *P* = 0.883; Table 5).

**Table 5.** Results of multiple linear regression models of *α* against length and number of genes within each SV. SV types: INV = inversion, DUP = duplication, INS = insertion, and DEL = deletion.

| SV type | Factor | Estimate | SE of estimate | *t*-value | *P*-value |
| --- | --- | --- | --- | --- | --- |
| DEL | length | 0.248 | 0.082 | 3.023 | **0.003** |
|  | number of genes | 0.016 | 0.110 | 0.148 | 0.883 |
|  | length: no. genes | -0.029 | 0.012 | -2.422 | **0.016** |
| DUP | length | 0.175 | 1.388 | 0.126 | 0.906 |
|  | number of genes | -0.841 | 1.132 | -0.743 | 0.499 |
|  | length: no. genes | 1.321 | 1.929 | 0.685 | 0.531 |
| INS | length | -0.171 | 0.520 | -0.328 | 0.745 |
|  | number of genes | -0.987 | 1.786 | -0.552 | 0.585 |
|  | length: no. genes | 0.105 | 0.135 | 0.779 | 0.443 |
| INV | length | -0.047 | 0.060 | -0.789 | 0.431 |
|  | number of genes | 0.185 | 0.079 | 2.348 | **0.021** |
|  | length: no. genes | -0.010 | 0.006 | -1.578 | 0.117 |

Finally, genetic differentiation (FST) between the *Lycaeides melissa* and Jackson Hole *Lycaeides* was moderately predictive of patterns of introgression in the hybrid zone. Specifically, F_ST_ was positively associated with the absolute value of α (Pearson *r* = 0.171, 95% CI = 0.120-0.222, *P* < 0.001) and negatively associated with signed values of α (Pearson *r* = −0.160, 95% CI: −0.210- -0.108, *P* < 0.001). Thus, SVs that differed more in frequency between the parental lineages exhibited more pronounced excess introgression overall, along with a specific shift towards excess Jackson Hole *Lycaeides* ancestry in the hybrid zone.

## Discussion

A comprehensive understanding of the genetic basis of speciation includes identifying the genes or mutations causing reproductive isolation and determining how often particular types of genetic change are involved (Serrato-Capuchina and Matute 2018). Hybrid zones provide a powerful tool to address both of these goals (Fitzpatrick et al. 2010; Harrison and Larson 2014; Gompert et al. 2017). First, quantifying variability in patterns of introgression across the genome in hybrid zones can lead to the identification of genomic regions or genes causing reproductive isolation in nature (Gompert and Buerkle 2009; Payseur 2010). Second, contrasting patterns of introgression across different categories of genetic or structural variants can help determine whether certain types of changes contribute disproportionately to speciation. Several models of speciation suggest a prominent role for SVs, including models of DMIs evolving following gene duplication, models involving recombination suppression via inversions, and various models of chromosomal speciation (e.g., Wright 1978; Lynch and Force 2000; Lynch et al. 2001; Feder and Nosil 2009; Zuellig and Sweigart 2018). Nonetheless, genome-wide characterization of patterns of structural variation needed to systematically address these hypotheses have been mostly lacking. We overcame this constraint using long-read Oxford nanopore sequencing and thereby quantified genome-wide patterns of introgression for SVs. We discuss technical aspects of our findings and the implications of our findings for understanding speciation below.

### Prevalence of SVs and sequencing technologies

We found that SVs are prevalent in *Lycaeides* butterflies, covering about 30% of the genome overall (though not 30% in any one individual). Considerably fewer SVs were detected using standard mate-pair sequencing (127,574 for nanopore vs 11,974 for mate-pair sequencing), but many that were found with mate-pair sequencing coincided with SVs from the nanopore data set (3891 out of 11974 SVs, 32%) (note that the two datasets also differed in the sampling scheme). This both bolsters our confidence in the nanopore data and highlights the utility of nanopore sequencing for SV genotyping relative to mate-pair sequencing. An additional 3842 SVs from the mate-pair data set were initially included in the nanopore data set but were removed during filtering because all individuals from one of the two parental species were heterozygotes. Most of the SVs with this pattern of excessive heterozygosity were deletions or insertions, consistent with expectations for pseudo-heterozygosity caused by errors in mapping transposable elements to the reference genome (Jaegle et al. 2021). The proportion of SVs exhibiting excessive heterozygosity from the mate-pair sequencing data set (~32%) was much higher than for the nanopore sequence data set (~12.7%). This discrepancy likely reflects the fact that short-read sequencing is unable to accurately capture the breakpoints of SVs that are located within repetitive regions of the genome. Thus, our results further highlight the advantages of applying long-read sequencing for identifying SVs.

### Structural variants, introgression and speciation

Consistent with past work (Chaturvedi et al., 2020), we found evidence that patterns of introgression varied across the genome, with some SVs showing excess Jackson Hole *Lycaeides* ancestry and some showing excess *L. melissa* ancestry. Moreover, across SVs there was an overall excess of loci with excess Jackson Hole *Lycaeides* ancestry relative to genome-average introgression based on SNP loci. Such a pattern would be expected if (i) selection in the hybrid zone, when it occurs, more frequently favors Jackson Hole *Lycaeides* alleles than *L. melissa* alleles, and if (ii) a greater proportion of SV loci are subject to selection than SNP loci, including direct and indirect selection caused by LD with actual barrier loci. Selection for Jackson Hole *Lycaeides* alleles could be driven by the environment, as the hybrid zone habitat is generally more like that found in nearby Jackson Hole populations, including a short summer season that is associated with univoltinism and obligate diapause as observed in *L. idas* and Jackson Hole *Lycaeides*, but not *L. melissa* (in contrast, the immediate agricultural habitat and host plant are indicative of *L. melissa)* (Gompert et al. 2013b). However, this result is also consistent with DMIs causing selection against minor-parent alleles (i.e., alleles from the parent species that has contributed less overall to an admixed population or hybrid zone) (Schumer et al. 2018; Martin et al. 2019; Chaturvedi et al. 2020). Lastly, because Jackson Hole *Lycaeides* are themselves ancient hybrids with *L. melissa* as one parent, their genomes might be enriched for alleles that function well in hybrids and thus are less likely to be selected against in a contemporary hybrid zone. Finally, the pattern that SVs show more evidence suggestive of selection than SNPs is consistent with recent studies of SVs in other organisms, including *Drosophila* (Weissensteiner et al. 2020) and grapevine (Zhou 2019).

Also consistent with past work (Chaturvedi et al., 2020), the Z sex chromosome was enriched for SV loci with patterns of introgression that deviated from the genome-wide average. This supports theory and empirical studies, which have found sex chromosomes plays a disproportional role in determining hybrid fitness (Coyne 1985; Masly and Presgraves 2007; Payseur and Rieseberg 2016; Gompert et al. 2017). The overall shift towards excess Jackson Hole *Lycaeides* ancestry noted above was also especially pronounced on the Z chromosome, which again could be explained by environment-dependent selection or an excess contribution to DMIs.

Several related hypotheses postulate a major role for inversions (one form of SV) in maintaining and promoting speciation (Noor et al. 2001; Rieseberg 2001; Hooper et al. 2019) (reviewed by Faria and Navarro 2010; Zhang et al. 2021a). We found little support for this hypothesis in *Lycaeides* butterflies Specifically, although inversions were more genetically differentiated between allopatric Jackson Hole *Lycaeides* and *L. melissa* populations, this increased differentiation did not result in exceptional patterns of introgression in the hybrid zone as would be expected if these loci were enriched for barrier loci. We did find an alternative, and somewhat unexpected pattern of introgression for inversions; signed estimates of α exhibited a modest, positive correlation with inversions containing more genes (Table 5, Figure 6B). In other words, in contrast to an overall signal of excess Jackson Hole ancestry in the hybrid zone relative to average ancestry based on SNPs, we detected excess *L. melissa* ancestry for inversions containing many genes. Because the hybrid zone has more individuals with Jackson Hole *Lycaeides* ancestry overall (Chaturvedi et al. 2020), this means that large inversions carrying excess *L. melissa* ancestry should frequently occur in a heterozygous state. One possible explanation for this pattern is that inversions with many genes contain a greater number of recessive deleterious mutations with different mutations associated with different inversion alleles, such that inversion heterozygotes would be heterozygous for these mutations and thus gain a fitness advantage via associative overdominance (Ohta 1971; Faria et al. 2019; Jay et al. 2021). Such an explanation was recently invoked to explain inversion polymorphisms among mimicry supergenes in *Heliconius* butterflies (Jay et al., 2021). More generally, empirical support from genomic studies for inversions contributing disproportionately to speciation has been mixed, with some studies finding no evidence that inversions play a major role in maintaining species boundaries, including sympatric *Heliconius* butterfly species (Davey et al. 2017) and *Timema* walking sticks (Lucek et al. 2019).

In contrast, deletions, which were the most common SV identified, exhibited the greatest overall deviations from genome-average patterns of introgression (507 of 562 SVs that deviated from null expectations of introgression were deletions). Moreover, ancestry informative deletions contain more genes than non-ancestry informative deletions (1000 permutation test: *p* <0.001; for a list the 169 annotated genes that are wholly or partially deleted by ancestry-informative deletions see Table S4). Some deletions might associate with phenotypic changes. For instance, one deletion showed excessive Jackson Hole *Lycaeides* ancestry contained gene *Fry* that could affect morphogenesis. Patterns of deletions are consistent with these loci being enriched for genetic variants contributing to reproductive isolation or those in linkage disequilibrium with such variants. Large deletions could generate underdominance of heterozygotes when potential functional genes are deleted or gene duplication followed by deletions, causing DMI (see table S4, Force et al. 1999; Lynch and Force 2000).

Finally, we found evidence that SVs that differed more between *L. melissa* and Jackson Hole *Lycaeides* exhibited excess directional introgression in Dubois, and specifically an excess of Jackson Hole *Lycaeides* ancestry. Still, genetic differentiation (F_ST_) only explained ~3% (squared correlation) of the variation in α. Particularly, inversions exhibit the highest F_ST_ between parental species compared to other SVs, but the lowest degree of deviation in introgression pattern. Taken together, these patterns highlighting the fact that differentiation in allopatry is far from a perfect predictor of introgression in hybrid zones and thus only somewhat informative of a genetic region’s contribution to speciation (Cruickshank and Hahn 2014; Burri 2017). This imperfect prediction of loci underlying genetic differentiation between *L. melissa* and Jackson Hole *Lycaeides* as barrier loci is consistent with previous study in hybrid zone between *L. melissa* and *L. idas* (Gompert et al. 2012a). The disconnection between loci underlying genetic differentiation and barrier loci could due to the limited genetic differentiation (F_ST_ = 0.05) where differentiated loci are less likely cause DMI among hybrids, or because of the environmental dependent hybrid fitness (Zhang et al. 2021b; Thompson et al. 2022). Altogether, our finding reveals the complexity of predicting the genetic basis of hybrid fitness based on parental differentiation.

## Acknowledgments

Funding for this project was provided by US NSF grants to ZG (1844941) and CCN (1050355). The support and resources from the Center for High Performance Computing at the University of Utah are gratefully acknowledged.

## Data Accessibility and Benefit-Sharing

Raw nanopore DNA sequence data will be deposited in the NCBI SRA (BioProject # pending). Mate-pair DNA sequence will be deposited in the NCBI SRA (BioProject # pending). Our new *L. melissa* genome assembly will be deposited in the NCBI Reference Sequence Database (accession number pending). Computer code and scripts central to this manuscript are available from GitHub: source code for genomic cline analysis (i.e., bgc) https://github.com/zgompert/BGC-Bayesian-genomic-clines, scripts for processing and analyzing the nanopore data (https://github.com/lz41/2022-Structural-variants), scripts for processing and analyzing the mate-pair DNA sequence data (https://github.com/lz41/2022-Structural-variants). Benefits from this research accrue from the sharing of our data and results on public databases as described above.

## Author Contributions

LZ and ZG designed the research. LZ, SC, CCN, LKL and ZG performed the research. LZ and SC analyzed the data. LZ wrote the paper. LZ, SC, CCN, LKL and ZG edited and revised the paper.

