## supplemental documents for "Population genomic evidence of selection on structural variants in a natural hybrid zone"

**Updated *L. melissa* genome using PacBio HiFi sequencing for gap filling**

We used PacBio SMRT sequencing to improve our existing *L. melissa* reference genome, which was originally assembled by Dovetail Genomics from Chicago and Hi-C genomic libraries and comprised 24 large scaffolds corresponding to the 23 *L. melissa* linkage groups (22 autosomes and the Z sex chromosome, with a all but one chromosome represented by a single scaffold) (Chaturvedi et al. 2020). Our primary goal was to reduce the proportion of the genome represented by Ns (i.e., unknown bases). High molecular weight DNA was extracted from a single female *L. melissa* by Brigham Young University (BYU) DNA Sequencing Center. Following quality control, a HiFi library was constructed from purified DNA and sequenced using a PacBio Sequell 2 with a 30-hour HiFi SMRT cell. This generated 978,737 (>Q20) reads with a mean length of 18,683 bps (total length = 18,286,022,199 >Q20 bps). The BYU DNA Sequencing Center bioinformatics team then used these data to improve (extend and replace Ns with known bases) our existing genome. Gaps were filled in the assembly from Dovetail Genomics using PacBio HiFi reads using TGS-GapCloser v1.0.1 (Xu *et al.* 2020). Because the reads were Q20 or higher, the "--ne" (i.e., skip error correction) option was used. Our input reference genome comprised 362 Mbps with 56% Ns, whereas after the gap-filling process our new reference genome spanned 521 Mbps with only 6.8% Ns.

We also updated genome annotation (i.e., gene locations and associated gene information) from our existing *L. melissa* genome (Chaturvedi et al. 2020) to the new version incorporating the PacBio sequence data. To do this, we first extracted the gene sequences from the original assembly based on the sequence coordinates (i.e., base pair positions)of the entire gene using samtools (version 1.9) (Li et al. 2009). We then performed a blast search to identify these same regions in the revised genome. This was done by first creating a local, blast database from the PacBio genome (using blastdb from blast 2.11.0) and then searching against this with each gene sequence. For this, we set the minimum e-value to 1e-50 and only considered alignments that were 92% or more identical. Then, using R, we identified the best hit for each gene sequence (where hits were ranked by match length weighted by percent identity with the target sequence), which we used to anchor the new gene coordinates. We used these new gene coordinates when performing analyses that included gene number or gene identity within the structural variant loci.

**Mate-pair DNA sequencing and structural variant calling**

As noted in the main text, we generated additional whole-genome sequence data to validate the SVs from the nanopore data. This data set included one butterfly from each of five nominal lineages of *Lycaeides* in North America: *L. idas* (from Trout Lake, WY), *L. melissa* (from Bonneville Shoreline Trail, UT), *L. anna* (from Yuba Gap, CA) and admixed lineages in the Sierra Nevada (from Carson Pass, CA) and Warner mountains (from Buck Mountain, CA) in the western USA. Genomic DNA was isolated from each butterfly using Qiagen's MatAttract HMW DNA extraction kit (Qiagen, Inc.), and a 3-kb mate-pair library was created for each butterfly species using the Nextera Mate-pair Library Gel-plus kit (Illumina, Inc.). The libraries were then sequenced on a HiSeq 2000 (2x100 bp reads) by Macrogen Inc. (Seoul, South Korea). We then aligned these data to the *L. melissa* reference genome using BWA MEM (version 0.7.17-r1188) (Li 2013). For alignment, we used a minimum seed length of 15 and only output alignments with a quality score of 30. We then used SAMTOOLS (version 1.12) (Li et al. 2009) to compress, sort, and index the alignments.

Both DELLY and LUMPY identify structural variants (SVs) based on breakpoint information relative to a reference genome. Both methods use the same information to identify SVs but apply different approaches. DELLY uses an integrative approach by combining shortrange and long-range paired-end mapping and split-read analysis to detect structural variation in the form of deletions, tandem duplications, inversions and translocations. Specifically, SVs are first identified based on disordantly mapped read pairs separated by unexpected distances given the insert size distribution; SV breakpoints are then localized from split-read alignments (Rausch et al. 2012, Lucek et al. 2019). On the other hand, LUMPY uses a probabilistic framework that integrates signals from multiple alignments to derive a general representation of a given SV breakpoint. This representation accounts for the variation in genomic resolution that is associated with different types of alignment evidence such as paired-end and split-read alignments (Layer et al. 2014)

We ran DELLY using sorted, indexed and duplicate-marked bam files for each of the five species included in the study. We ran the DELLY CALL function by specifying standard parameters as follows: the minimum paired-end (PE) mapping quality (-q) was set to 1, the minimum PE quality for translocation (-r) was set to 20, the insert size cutoff (-s) was set to 9, the minimum clipping length (-c) was set to 25, the maximum read separation (-n) was set to 40, and the minimum mapping quality for genotyping (-u) was set to 5. We converted the BCF output from DELLY CALL command to VCF format, using BCFTOOLS (version 1.6). We then merged the calls from individual species to get a final merged VCF for all species using the DELLY MERGE command.

Before identifying SVs using LUMPY, we estimated the empirical insert sizes for each of the five species using SAMTOOLS view and the pairedend_distro.py script from LUMPY. Based on these estimates, we ran LUMPY by setting the mean and standard deviations (SDs) of paired-end insert sizes for each species as follows: *L. anna*: mean = 327, SD = 172; Sierra Nevada: mean = 337, SD=179; *L. idas:* mean = 305, SD = 169; Warner mountains: mean = 324, SD = 175, and

*L. melissa*: mean = 305, SD = 173. Additionally, we specified the following conditions for identifying SVs: the minimum weight (-weight) for a call was set to 4, the minimal mapping quality (-min_mapping_threshold) was set to 20, a window of ±10 bp around a split alignment was included in each breakpoint interval (-back_distance), and the number of unique bases at the end of each read pair was required to be equal to the read length (101 bps, -read_length) so that paired reads would not overlap.

**Table S1.** Summary of sample collection information. Latitude and longitude are reported in degrees.

| **Species** | **Locality** | **ID** | **Longitude** | **Latitude** | **Number of individuals** |
|---|---|---|---|---|---|
| Jackson hole hybrids | Frontier Creek, WY | FRC | -109.5776 | 43.7206 | 4 |
| Jackson hole hybrids | Mt. Randolf, WY | MRF | -110.3918 | 43.8547 | 3 |
| Dubois hybrids | Dubois, WY | DBS | -109.6991 | 43.5623 | 26 |
| *L. melissa* | Sinclair, WY | SIN | -107.1131 | 41.8511 | 4 |
| *L. melissa* | Victor, ID | VIC | -111.1114 | 43.659 | 2 |

**Table S2.** Sequencing libraries for each individual

| Collection year | Sample ID | Sample in Chaturvedi et al. 2020 | DNA concentration (ng/ul) | Multiplexing library |
|---|---|---|---|---|
| 2016 | DBS-16-07 | YES | 3.02 | HL1 |
| 2016 | DBS-16-16 | YES | 8 | HL1 |
| 2016 | DBS-16-22 | YES | 18.8 | HL1 |
| 2020 | SIN-20-01 | NO | 4.18 | HL1 |
| 2013 | DBS-13-04 | YES | 3.86 | HL2 |
| 2014 | DBS-14-02 | YES | 3.22 | HL2 |
| 2016 | DBS-16-15 | YES | 3.04 | HL2 |
| 2016 | DBS-16-20 | YES | 2.22 | HL2 |
| 2016 | DBS-16-28 | YES | 2.96 | HL2 |
| 2020 | MRF-20-04 | NO | 3.86 | HL2 |
| 2020 | SIN-20-04 | NO | 5.26 | HL2 |
| 2020 | SIN-20-05 | NO | 4.48 | HL2 |
| 2018 | VIC-18-12 | NO | 4.06 | HL2 |
| 2013 | DBS-13-08 | YES | 2.14 | HL2, HL6 |
| 2014 | DBS-14-06 | YES | 2.14 | HL2, HL6 |
| 2014 | DBS-14-08 | YES | 1.85 | HL2, HL6 |
| 2013 | DBS-13-09 | YES | 5.6 | HL3 |
| 2016 | DBS-16-14 | YES | 10 | HL3 |
| 2016 | DBS-16-18 | YES | 6.18 | HL3 |
| 2016 | DBS-16-21 | YES | 9.64 | HL3 |
| 2018 | VIC-18-14 | NO | 7.44 | HL3 |
| 2018 | VIC-18-15 | NO | 5.64 | HL3 |
| 2016 | DBS-16-10 | YES | 12.7 | HL4 |
| 2016 | FRC-16-01 | NO | 11.3 | HL4 |

| | | | | |
|---|---|---|---|---|
| 2016 | FRC-16-16 | NO | 11.8 | HL4 |
| 2020 | MRF-20-01 | NO | 11.2 | HL4 |
| 2020 | MRF-20-03 | NO | 12.1 | HL4 |
| 2017 | VIC-17-01 | NO | 14.1 | HL4 |
| 2014 | DBS-14-17 | YES | 15.5 | HL5 |
| 2016 | DBS-16-08 | YES | 21.2 | HL5 |
| 2016 | DBS-16-19 | YES | 25.4 | HL5 |
| 2016 | DBS-16-23 | YES | 21.2 | HL5 |
| 2016 | DBS-16-24 | YES | 23 | HL5 |
| 2016 | DBS-16-27 | YES | 19.5 | HL5 |
| 2016 | FRC-16-12 | NO | 48 | HL6 |
| 2016 | FRC-16-15 | NO | 27 | HL6 |
| 2014 | DBS-14-15 | YES | NA | HL7 |
| 2016 | DBS-16-26 | YES | 3.6 | HL8 |
| 2020 | SIN-20-02 | NO | 4.54 | HL9 |

**Table S3.** Results of multiple linear regression models of the absolute value of $\alpha$ against SV length and number of genes within each SV. SV types: INV = inversion, DUP = duplication, INS = insertion, and DEL = deletion.

| **SV types** | **Factor** | **Estimate** | **SE of estimate** | ***t*-value** | ***P*-value** |
|---|---|---|---|---|---|
| DEL | length | -0.033 | 0.038 | -0.866 | 0.387 |
| | number of genes | 0.024 | 0.046 | 0.523 | 0.601 |
| | length: no. genes | -0.001 | 0.005 | -0.109 | 0.913 |
| DUP | length | 0.037 | 0.309 | 0.12 | 0.91 |
| | number of genes | 0.138 | 0.137 | 1.005 | 0.372 |
| | length: no. genes | -0.755 | 0.356 | -2.123 | 0.101 |
| INS | length | -0.031 | 0.173 | -0.18 | 0.858 |
| | number of genes | -0.001 | 0.259 | -0.004 | 0.997 |
| | length: no. genes | 0.004 | 0.036 | 0.11 | 0.913 |
| INV | length | -0.007 | 0.037 | -0.199 | 0.843 |
| | number of genes | 0.048 | 0.042 | 1.152 | 0.252 |
| | length: no. genes | -0.004 | 0.003 | -1.327 | 0.187 |

Note: estimates are standardized regression coefficients.

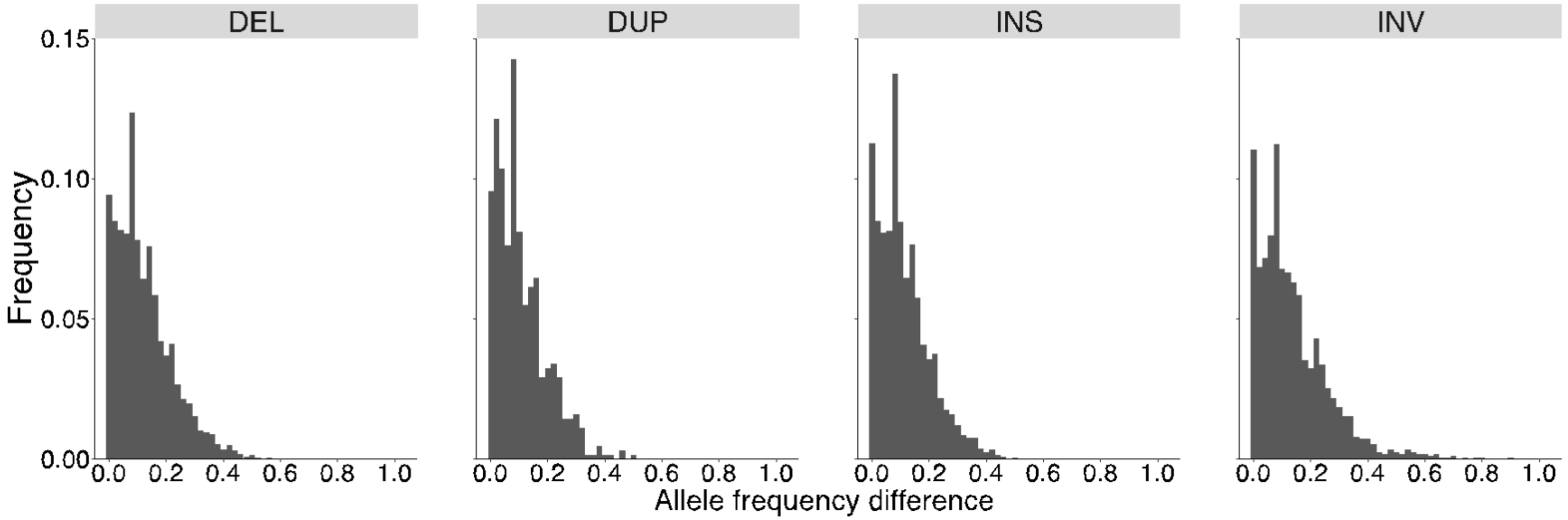


**Figure S1.** Histograms of distribution of allele frequency differences between Jackson Hole *Lycaeides* and *L. melissa* for different types of structural variants.

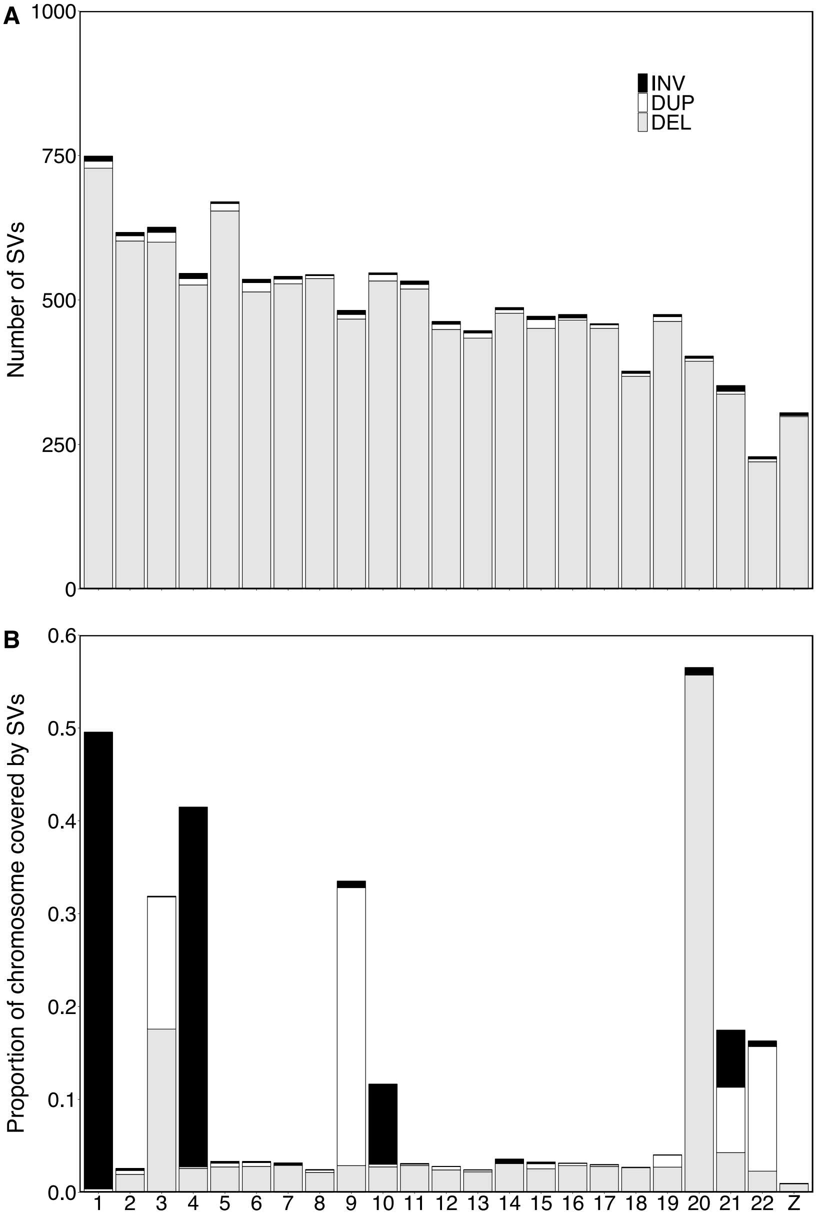


**Figure S2.** (A) Number of SVs identified by mate-pair data across the genome. (B) Proportion of each chromosome covered by SVs identified by mate-pair data. A few chromosomes have extremely high coverage because of a few very large SVs were detected on these chromosomes.

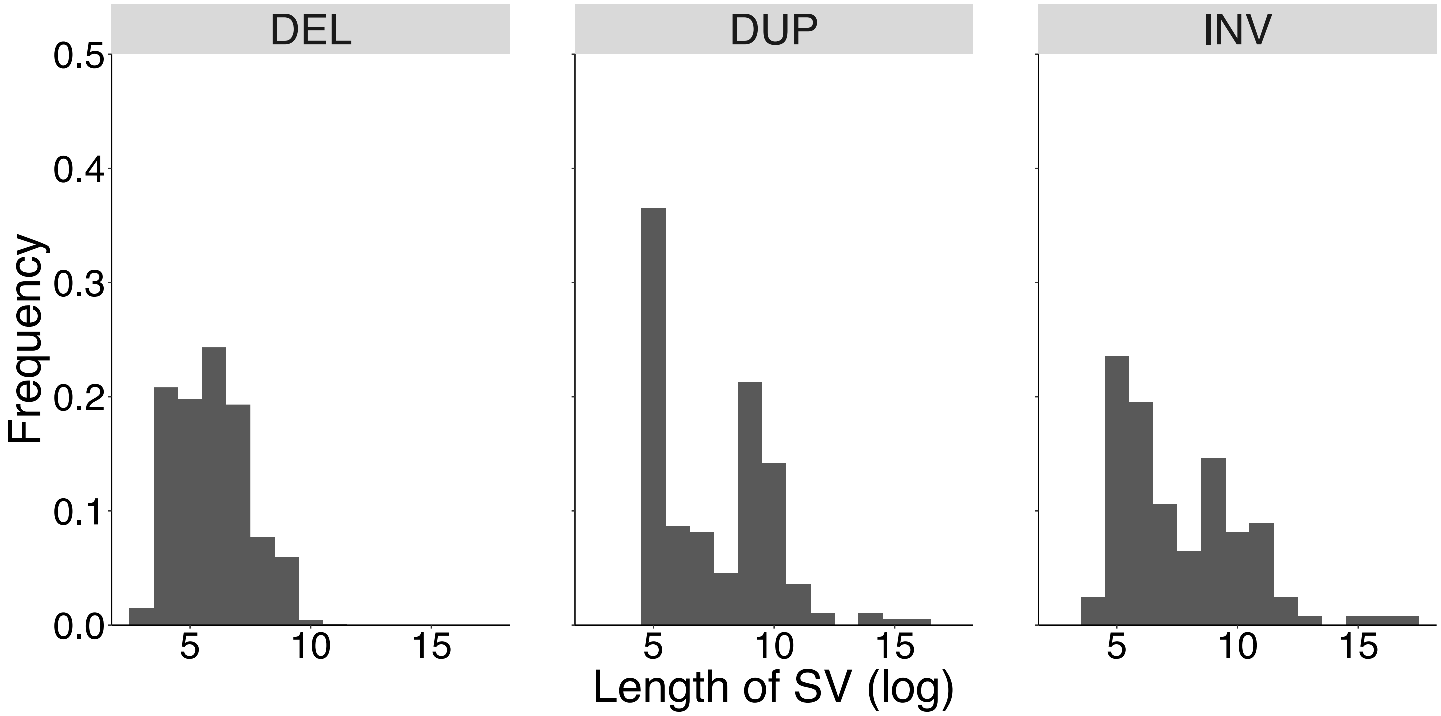


**Figure S3.** Histogram showing the distribution of length of SVs (log 10) identified by mate-pair data

**Results of SVs detected with $\varepsilon$ = 0.05, $\varepsilon$ stands for the probability of a sequence read erroneously supporting the reference or structural variant allele.**

***Number, length, and distribution of SVs across genome***

A total of 86,788 SVs were identified that covered 21.5% of the genome, including 11,199 SVs >1000 bp in length. Among all SVs identified, there were 50,503 deletions covering 13.9% of the genome, 34,964 insertions covering 2.2% of the genome, 985 inversions covering 4.9% of the genome and 336 duplications covering 0.3% of the genome (see Table S5 for range of SV sizes; Figure S4A, 4B). Considering only SVs > 1kbp in length, we identified 10,052 deletions covering 12.1% of the genome, 717 inversions covering 4.8% of the genome, 365 insertions covering 0.8% of the genome and 65 duplications covering 0.3% of the genome (see Table S5 for range of SV sizes, Figure S4C, D).

***Population genomic patterns among structural variants***

A total of 1403 SVs exhibited allele frequency differences exceeding 0.3 (compared to N = 1419 for error rate = 0.01). Population differentiation measured by SVs $F_{ST}$= 0.054, was generally similar to earlier estimates for SNPs ($F_{ST}$ = 0.05; see Chaturvedi et al. 2020) and estimates for SVs with error rate= 0.01 ($F_{ST}$ = 0.048) (Table S6, Figure S3B, 3C)). However, genetic differentiation ($F_{ST}$) for inversions was higher than for other SV types or SNPs (Table S6).

***Patterns of introgression for SVs in the Dubois hybrid zone***

Credible variation in patterns of introgression across the genome relative to genome-average ancestry based on SNPs were detected (i.e., cases where 95% equal-tail probability intervals [ETPIs] for genomic cline parameters did not overlap with zero) (Figure S4B, C). A total of 122 SVs showed credible excess introgression from *L. melissa* (95% ETPIs for cline parameter $\alpha$ > 0) and 293 SVs exhibited credible excess introgression from Jackson Hole *Lycaeides* (95%

ETPIs for cline parameter $\alpha < 0$). Consequently, relative to the average introgression for SNPs, SVs showed an excess of Jackson Hole *Lycaeides* ancestry in the hybrid zone (mean $\alpha \pm$ SE = -0.205 ± 0.017; probability of getting 293 credible excess introgress loci out of 415 loci with cline parameter $\alpha < 0$: $P < 0.0001$, Figure S4C). No SVs displayed credible shifts in the rate or extent of introgression (95% ETPIs for cline parameter $\beta > 0$ or $\beta < 0$). Estimates of α based on assumed error rates ($\varepsilon$) of 0.05 or 0.01 were highly correlated ($r = 0.995$, $P < 0.0001$), and thus quantitative patterns of introgression were robust to the different assumed error rates.

Similar to the results shown in the main text, SVs with excess directional introgression relative to genome-average ancestry (based on SNPs) were distributed across the 23 *Lycaeides* chromosomes, but SVs on Z-chromosome showed significantly more excess Jackson Hole *Lycaeides* introgression compared to SVs on the autosomes (permutation test with 1000 permutations: mean difference in $\alpha$, [Z - autosomes] = -0.315, $P < 0.001$).

**Table S5.** Number and length (mean, 2.5th percentile and 97.5th percentile) of SVs with $\varepsilon = 0.05$ detected by nanopore sequencing. SV types: INV = inversion, DUP = duplication, INS = insertion, and DEL = deletion.

| | SV type | No. SVs | Length of SVs (bp) | | |
|---|---|---|---|---|---|
| | | | Mean | 2.50% | 97.50% |
| **All SVs** | DEL | 50503 | 1586 | 42 | 11532 |
| | DUP | 336 | 4951 | 51 | 22088 |
| | INS | 34964 | 328 | 42 | 702 |
| | INV | 985 | 39965 | 186 | 381392 |
| **SVs with length>1000 bp** | DEL | 10052 | 7037 | 1034 | 27946 |
| | DUP | 65 | 24297 | 1050 | 68163 |
| | INS | 365 | 11553 | 1000 | 65049 |
| | INV | 717 | 54727 | 1119 | 488219 |

**Table S6.** Comparison of pairwise $F_{ST}$ across different structural variant types with $\varepsilon$ = 0.05 detected by nanopore sequencing. Lineage codes: JH = Jackson Hole *Lycaeides*, LM = *L. melissa*, DBS = Dubois hybrid zone. SV types: INV = inversion, DUP = duplication, INS = insertion, and DEL = deletion.

| **Pairs** | **Fst (all SVs)** | **Fst (INV)** | **Fst (DUP)** | **Fst (INS)** | **Fst (DEL)** |
|---|---|---|---|---|---|
| JH-LM | 0.054 | 0.09 | 0.051 | 0.051 | 0.055 |
| DBS-JH | 0.026 | 0.047 | 0.028 | 0.024 | 0.026 |
| DBS-LM | 0.035 | 0.058 | 0.031 | 0.036 | 0.034 |

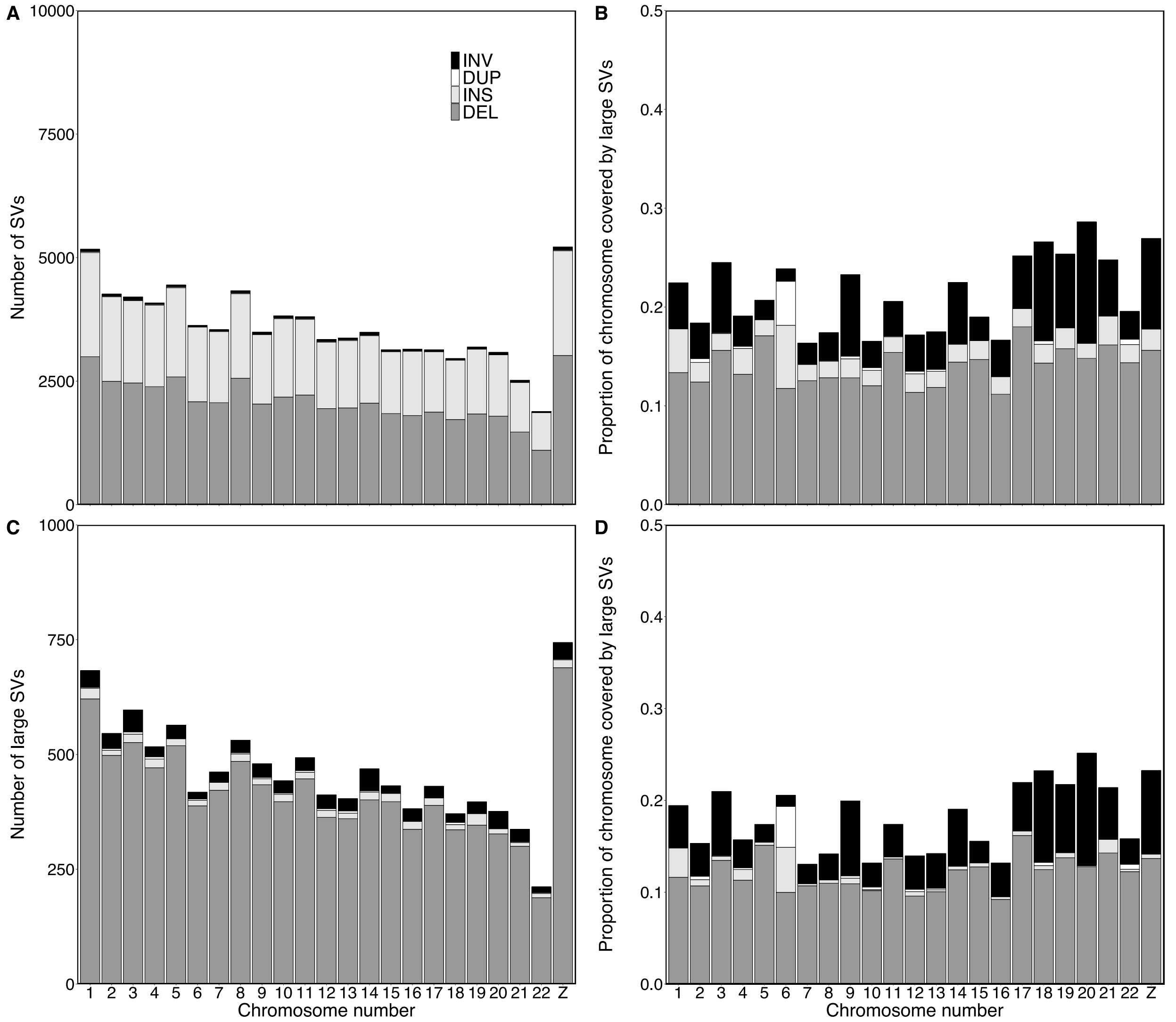


**Figure S4.** (A) Number of SVs with $\varepsilon = 0.05$ detected by nanopore sequencing across the genome. (B) Proportion of each chromosome covered by SVs. (C) Number of large SVs (>1000 bp) across the genome. (D) Proportion of each chromosome covered by large SVs (>1000 bp). Bars with different colors denote different SV types, as shown in the legend: INV = inversion, DUP = duplication, INS = insertion, and DEL = deletion.

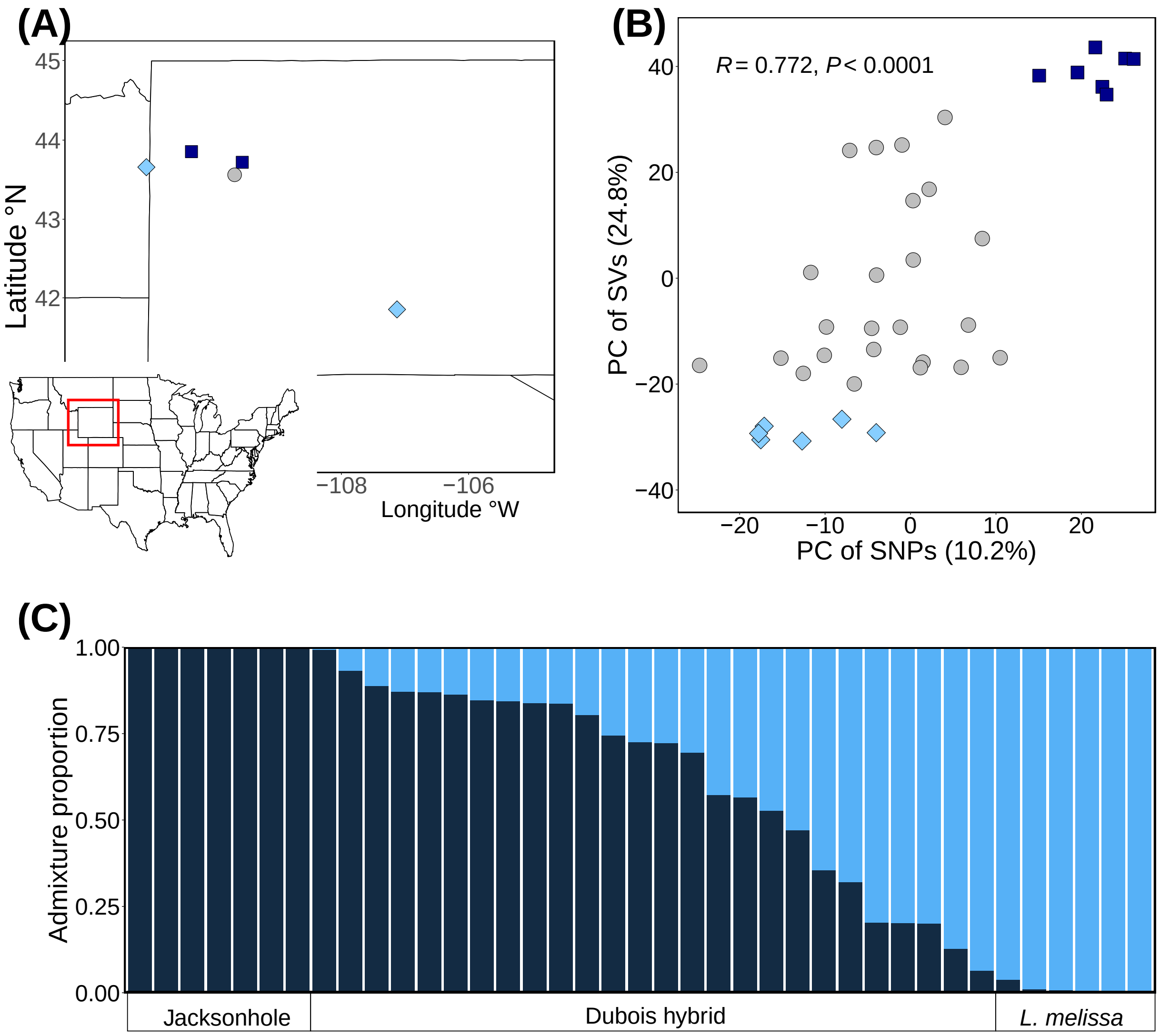


**Figure S5.** (A) Collection sites, (B) Plot of PC 1 scores result for large, ancestry informative SVs with $\varepsilon$ = 0.05 (>1000bp, allele frequency differences>0.3) vs. PC2 scores of SNPs, (C) Entropy plot from large, ancestry informative SVs ($\varepsilon$ = 0.05, >1000bp, allele frequency differences>0.3), y axis is the Bayesian estimates of admixture proportion.

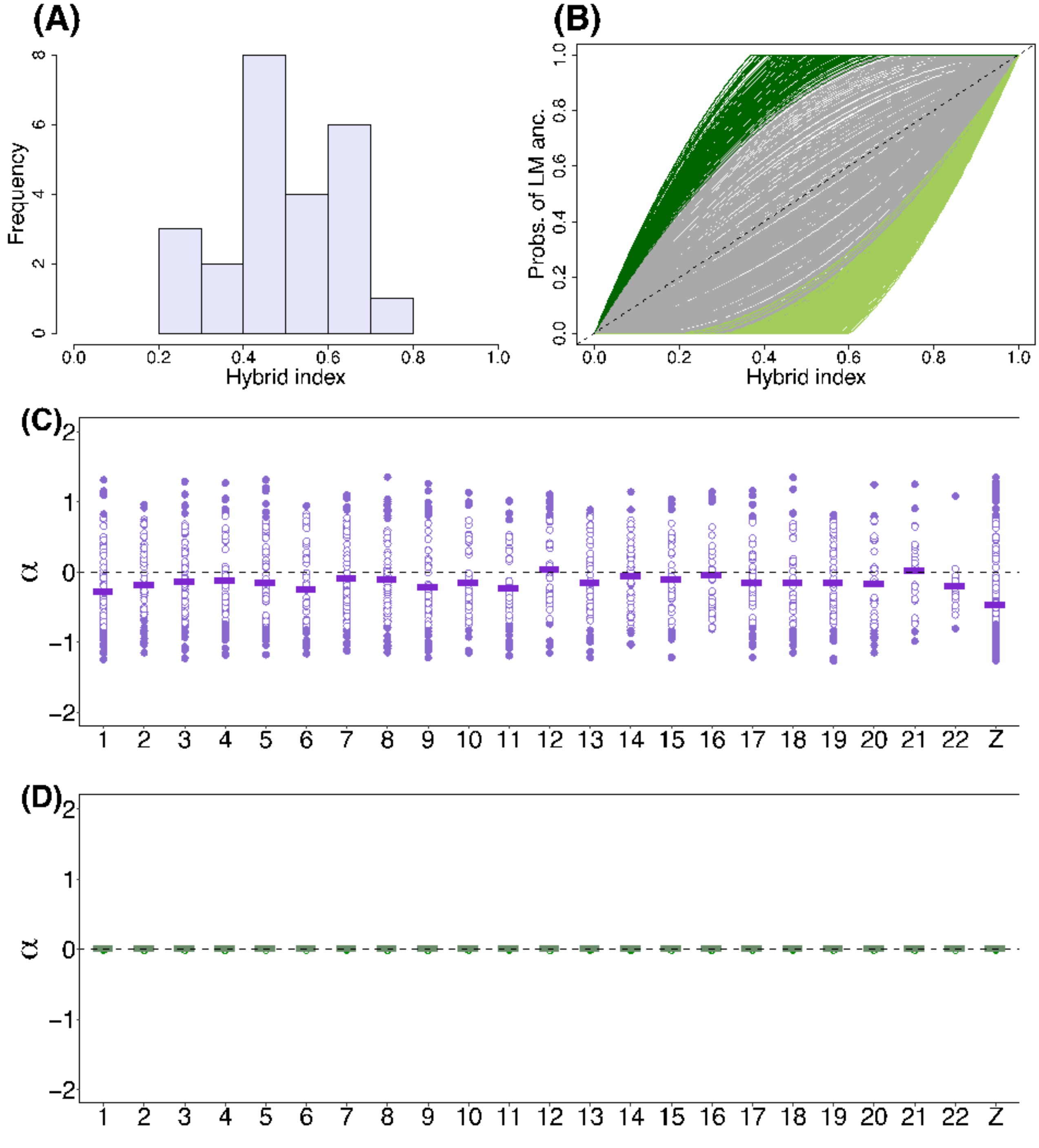


**Figure S6.** Summary of the genomic cline analysis of SVs with $\varepsilon = 0.05$. (A) Distribution of hybrid index using structural variant data. (B) Estimated genomic cline relative to SNPs using ancestry-informative SVs. Each solid line gives the estimated probability of Jackson Hole (JH) ancestry for a structural variant locus. Green lines denote cases of credible directional introgression (95%

ETPIs for α that do not overlap with zero), gray lines denote clines not significantly different from the genome average. The dashed line gives the null expectation based on genome-wide admixture. Distribution of cline parameters $\alpha$ (**C**) and $\beta$ (**D**) across different chromosomes. Solid circles indicate loci with cline parameters indicating credible deviations from genome-average introgression (95% ETPIs that do not overlap with zero), whereas unfilled circles indicate loci not significantly different from the genome average. Solid horizontal lines are the mean value of cline parameters across all loci at each chromosome. The dashed horizontal line gives the null expectation based on genome-wide admixture using SNP loci.

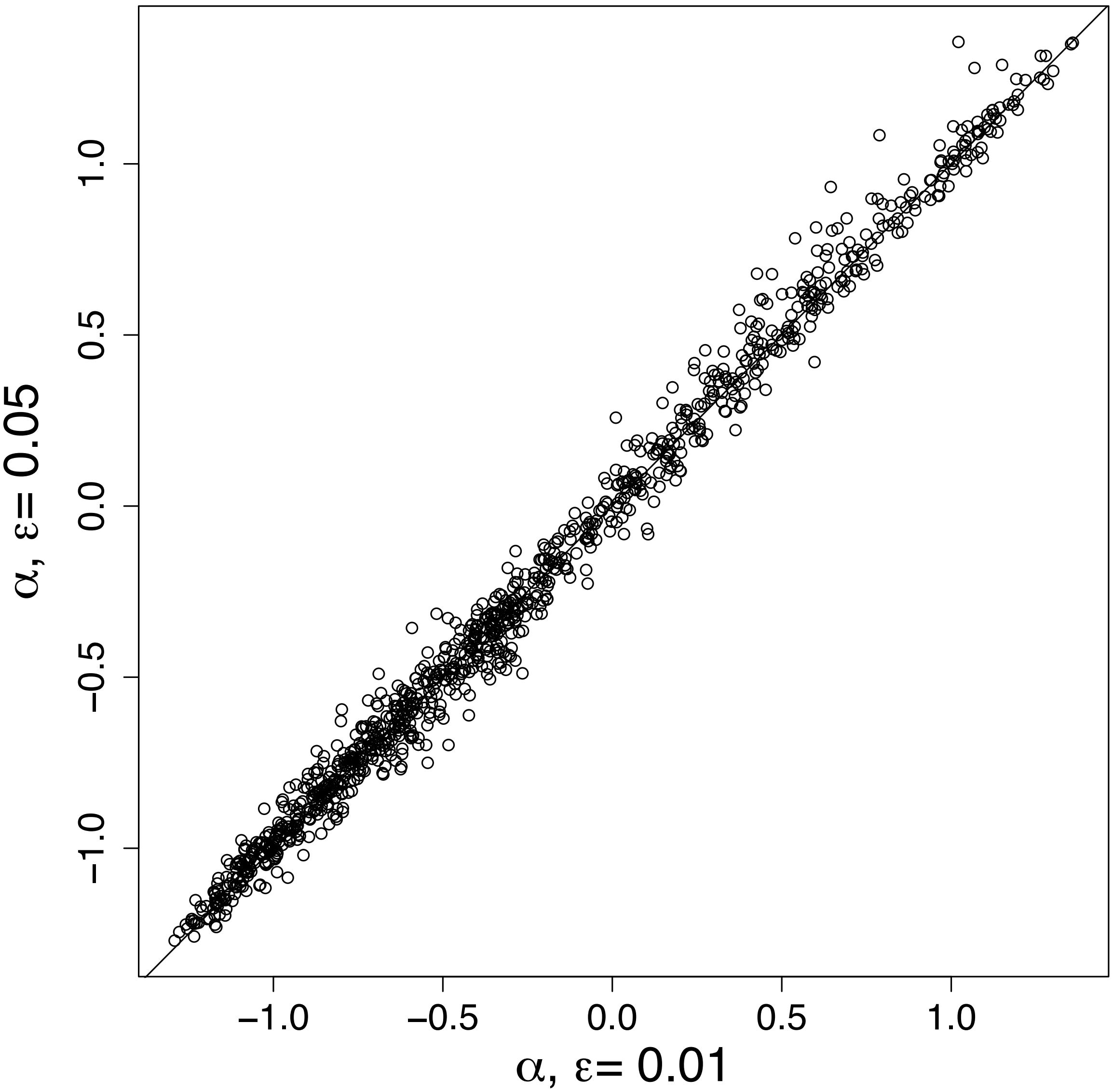


**Figure S7.** Scatterplot showing the correlation between point estimates (posterior means) of $\alpha$ for SVs with error rates ($\varepsilon$) of 0.01 versus 0.05.
